# Allosteric modulation by the fatty acid site in the glycosylated SARS-CoV-2 spike

**DOI:** 10.1101/2023.11.06.565757

**Authors:** A. Sofia F. Oliveira, Fiona L. Kearns, Mia A. Rosenfeld, Lorenzo Casalino, Lorenzo Tulli, Imre Berger, Christiane Schaffitzel, Andrew D. Davidson, Rommie E. Amaro, Adrian J. Mulholland

## Abstract

The trimeric spike protein plays an essential role in the SARS-CoV-2 virus lifecycle, facilitating virus entry through binding to the cellular receptor angiotensin-converting enzyme 2 (ACE2) and mediating viral-host membrane fusion. The SARS-CoV-2 spike contains a fatty acid (FA) binding site at the interface between two neighbouring receptor-binding domains. This site, also found in some other coronaviruses, binds free fatty acids such as linoleic acid. Binding at this site locks the spike in a non-infectious, closed conformation. This site is coupled to functionally important regions, but the effects of glycans on these allosteric effects have not been investigated. Understanding allostery and how this site modulates the behaviour of the spike protein could potentiate the development of promising alternative strategies for new coronavirus therapies. Here, we apply dynamical nonequilibrium molecular dynamics (D-NEMD) simulations to investigate allosteric effects of the FA site in the fully glycosylated spike of the original SARS-CoV-2 ancestral variant. The results show allosteric networks that connect the FA site to important functional regions of the protein, including some more than 40 Å away, including the receptor binding motif, an antigenic supersite in the N-terminal domain, the furin cleavage site, regions surrounding the fusion peptide, and another allosteric site known to bind heme and biliverdin. The networks identified here highlight the complexity of the allosteric modulation in this protein and reveal a striking and unexpected connection between different allosteric sites. Notably, 65% of amino acid substitutions, deletions and insertions in the Alpha, Beta, Delta, Gamma and Omicron variants map onto or close to the identified allosteric pathways. Comparison of the FA site connections from D-NEMD in the glycosylated and non-glycosylated spikes revealed that the presence of glycans does not qualitatively change the internal allosteric pathways within the protein, with some glycans facilitating the transmission of the structural changes within and between subunits.

**Significance statement:** The spike protein is crucial for the SARS-CoV-2 virus, enabling the fusion of the viral and host cell membranes. This protein contains several allosteric sites, including a fatty acid binding site at the interface between every two neighbouring receptor-binding domains. This site modulates the behaviour of the protein, with the binding of various free fatty acids and other small molecules influencing the spike’s structure. In particular, the binding of linoleic acid, an essential fatty acid molecule, stabilizes the protein in a non-infectious locked conformation, thus making it inaccessible for binding to human receptors. Here, we investigate how the fatty acid site modulates the structural and dynamical behaviour of the fully glycosylated protein. Our work reveals complex patterns of communication between the fatty acid site and functionally important regions of the spike (including the receptor binding motif, the antigenic supersite in the N-terminal domain, the heme/biliverdin site, furin cleavage site and the fusion-peptide surrounding regions) and shed new light on the roles of glycans in this protein.

## Main text

The SARS-CoV-2 virus, like other β-coronaviruses, uses the spike protein to mediate virus entry into host cells. The spike is a trimeric glycoprotein embedded in the virus envelope. During the initial stage of the SARS-CoV-2 infection process, the spike binds to host target cells, primarily *via* the receptor angiotensin-converting enzyme 2 (ACE2) (1, 2), but it can also bind to other targets, such as neuropilin-1 (3, 4), estrogen receptor α (5) and potentially to nicotinic acetylcholine receptors (6–9) and sugar receptors (10). Given its crucial role in the infection process and the fact that it is one of the main targets for antibody neutralization, the spike is one of the most important targets for developing COVID-19 therapies and vaccines (e.g. (11–15)).

Each spike monomer comprises three regions: a large ectodomain, a transmembrane domain and a short cytoplasmic tail (16–18). The ectodomain, the main focus of this work, is composed of two subunits (S1 and S2) and contains the structural motifs that directly bind to the host receptors as well as those needed for the membrane fusion process (16–18). S1 is responsible for binding to the human ACE2 receptors, while S2 for the fusion of the viral and host membranes (16–18). The spike also contains three equivalent fatty acid (FA) binding sites at the interfaces between neighbouring receptor binding domains (RBDs) (19) (Figure 1A). Each FA binding site is a hydrophobic pocket formed by two RBDs, with one RBD providing the aromatic and hydrophobic residues to accommodate the FA hydrocarbon tail and the other providing the polar and positively charged residues that bind the FA carboxylate headgroup (Figure 1B). The essential FA linoleic acid binds (as linoleate, LA) with high affinity to the FA pocket, stabilizing the spike in a non-infectious locked conformation, in which the RBDs are all ‘down’ with the receptor-binding motifs (RBMs) occluded inside the trimer, and thus inaccessible for binding to ACE2 (19). The discovery of this site inspired the development of new spike-based potential therapies based on FAs or other natural, repurposed, or specifically designed small molecules able to bind to the FA site (11, 20–24). Several cryo-EM structures of the SARS-CoV-2 spike in complex with small molecules, such as linoleic, oleic, and all-trans retinoic acid and SPC-14, bound to the FA site, are now available (11, 19–21). Following this discovery, equivalent FA sites have been identified in several closely related coronavirus spikes (19, 25, 26). Surface plasmon resonance experiments and cryo-EM structures show that the FA site is conserved in the spike proteins of highly several pathogenic β-coronaviruses, such as SARS-CoV, MERS-CoV, SARS-CoV-2, but not in the spikes of common, mild disease-causing β-coronaviruses (27).

**Figure 1.**
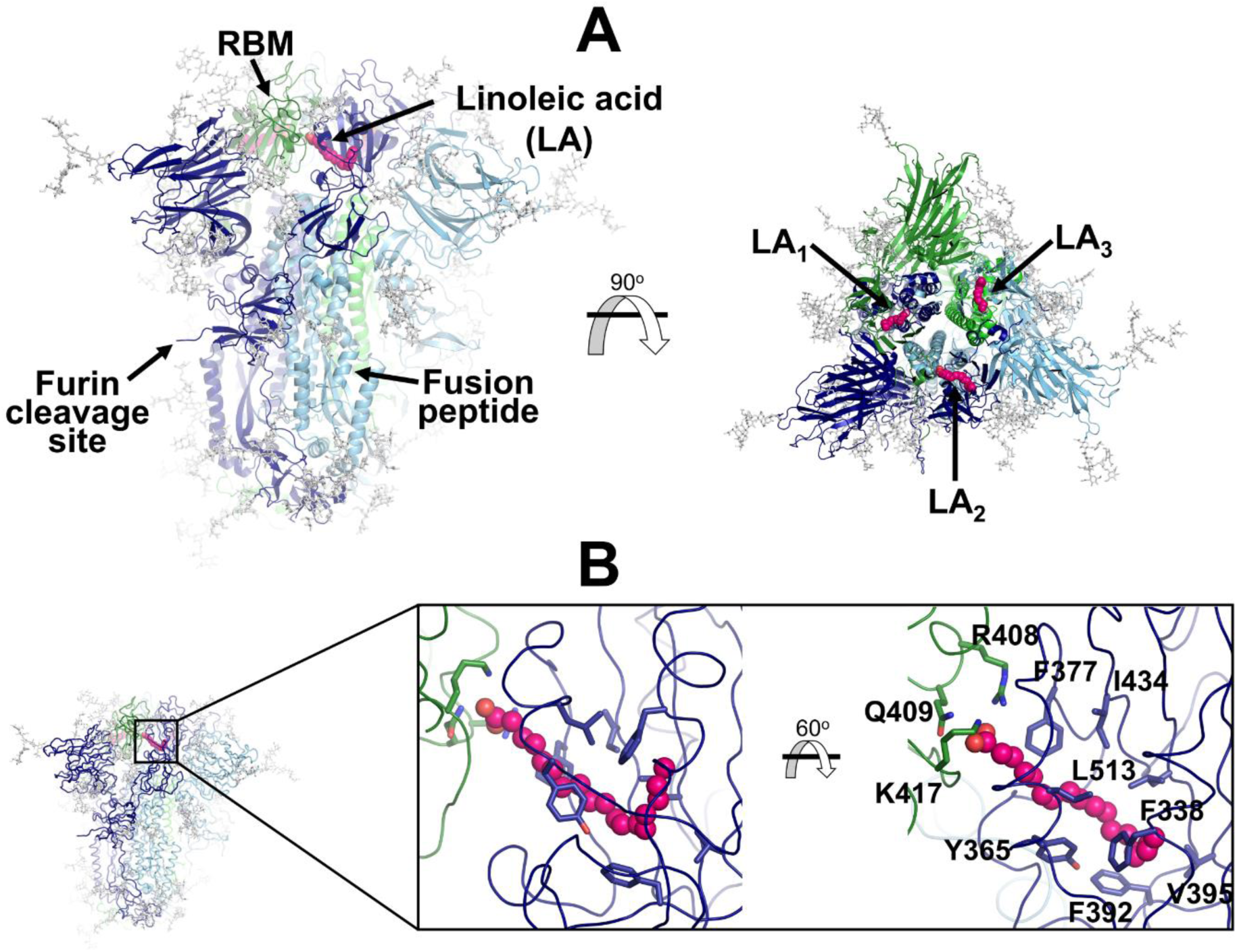
Structure of the glycosylated head region of the ancestral SARS-CoV-2 spike with linoleate bound to the free fatty acid (FA) binding site. (A) Model of the ectodomain of the glycosylated SARS-CoV-2 spike with linoleate (LA) bound. The spike-LA complex model was built using the cryo-EM structure 7JJI as a reference (26). Each monomer in the spike homotrimer is shown in a different colour: dark blue, light blue and green. Glycans are indicated with grey sticks, and LA molecules are highlighted with magenta spheres. Three FA binding sites exist in the trimer, each located at the interface between two neighbouring monomers. In this model, all three receptor-binding motifs (RBMs) are in the ‘down’ conformation, and the protein is cleaved at the furin recognition site at the S1/S2 interface. (B) Detailed view of the FA binding site. This hydrophobic site is formed by two RBDs, with one providing the hydrophobic pocket for the FA hydrocarbon tail and the other providing polar (Q409) and positively charged (R408 and K417) residues to bind the negatively charged FA headgroup.

Simulations using the dynamical-nonequilibrium molecular dynamics (D-NEMD) approach showed that the FA site allosterically modulates the behaviour of functional motifs in both the ancestral (also known as wild type, ‘early 2020’, or original) spike and in several variants (28–30), and that these effects differ between variants. These D-NEMD simulations (which tested the effects of removing linoleate) showed that the FA site is allosterically connected to the RBM, N-terminal domain (NTD), furin cleavage site, and the region surrounding the fusion peptide (FP) (28–30). These regions have significantly different allosteric behaviours between the ancestral, Alpha, Delta, Delta plus, and Omicron BA.1 variants (28–30). They differ not only in the amplitude of the structural responses of these regions but also in the rates at which the structural changes propagate (28–30). However, these previous D-NEMD simulations did not incorporate the spike’s many *N-* and *O-*linked glycans, which are critical to its function, not only in protecting it from immune recognition, but also in modulating its dynamics (31–34). A crucial unresolved question, therefore, is whether and how glycosylation affects allosteric communication with the FA site.

The ancestral spike is heavily glycosylated with 22 predicted *N*-linked glycosylation sites per monomer (16, 35), of which 17 have been found to be occupied (35–38). The ancestral spike also contains at least two *O-*glycosylation sites per monomer with low occupancy (37, 39, 40). The occupancy and composition profile of the glycosylation sites differ between variants, expression systems, and experimental methods (41, 42). This glycan coating plays a crucial role in shielding the virus from the immune system (33, 34), and in infection (31–34, 43). Glycans also affect the dynamics and stability of essential regions of the protein, including the RBDs, and modulate binding to ACE2 (31–33). Here, we use D-NEMD simulations (44–46) to characterize the response of the fully glycosylated SARS-CoV-2 ancestral spike to LA removal, and investigate allosteric modulation by the FA site and the effects of glycans on the protein’s allosteric behaviour. In recent years, D-NEMD simulations (44–46) have emerged as a powerful computational approach to investigate a diversity of biological problems from the trasmission of structural changes (47, 48) to the identification of allosteric effects (28–30, 49–55) and the impact of pH changes (56) in fundamentally different biomolecular systems, including SARS-CoV-2 targets (28–30, 54, 56). For example, in the SARS-CoV-2 main protease, dynamical responses from D-NEMD pinpointed positions associated with drug resistance (54), and for the spike, D-NEMD indicated the regions of the protein affected by pH changes (56).

We performed extensive equilibrium MD simulations, followed by hundreds of D-NEMD simulations, to analyze the response of the fully glycosylated, cleaved (at the furin recognition site) spike to LA (linoleate) removal. Principal component analysis was performed to check the equilibration and sampling of the equilibrium replicates (Figure S1C). In the equilibrium simulations, the locked state of the spike (with all RBDs down) with linoleate bound remained stable (Figure S1A), showing structural convergence after ∼50 ns and minimal secondary structure loss after 750 ns (Figure S1B). The comparison between the average C_α_ fluctuations calculated from the equilibrium simulations for the locked (this work) and closed (from Casalino *et al.* (33)) glycosylated spikes shows that the dynamics of the protein is generally similar (Figure S1D). The largest difference is observed in RBM_B_, which exhibits decreased dynamics in the locked state (i.e. when LA is present in the FA sites) (Figure S1D). In the locked spike equilibrium simulations, all LA molecules remained stably bound to the protein (Figures S2A and S2D), with the carboxylate head group of LA making consistent salt-bridge interactions with K417 and occasional interactions with R408 (Figures S2B and S2C).

Analysis of the dynamics of the glycans in the equilibrium trajectories (Figures S3-S4) showed (as previously observed for the spike without LA (33)) that the glycans are very mobile, exhibiting diverse levels of motion depending on their composition, branching and solvent exposure (Figure S3). Generally, *N*-glycans in the NTD show higher fluctuations than those of the RBD (Figure S3). The glycan linked to N331 from chain C is an exception, with one of the largest RMSF values. The *O-*glycans connected to T323 and S325, close to the RBD, are less flexible than the *N*-glycans (Figure S3). The highly dynamic profile of the glycans (Figure S4) helps the spike to evade the host immune response by masking immunogenic epitopes, thus preventing them from being targeted by the host’s neutralizing antibodies. To quantify the shielding effect, the spike accessible surface area (ASA) covered by the glycans was determined for probe radii ranging from 0.14 nm (approximate radius of a water molecule) to 1.1 nm (approximate radius of a small antibody molecule). As can be seen in Figures S4B and S4C, consistently with previous findings reported for the closed state (with all RBDs in the ‘down’ conformation, without LA bound) (33), in the locked state, the spike head has a thick glycan shield, which covers ∼60% of the protein accessible area for a 1.0-nm-radius probe and restricts the binding of medium size molecules to the protein. However, small molecules (probes with a radius 0.14-0.3nm) can penetrate the shield more easily as it only covers ∼26% of the area of the protein accessible to smaller probes (Figure S4C).

An ensemble of 210 conformations (70 configurations per replicate) was extracted from the equilibrium MD simulations and used as starting points for the D-NEMD simulations, which investigated the effect of LA removal (Figure S6). The D-NEMD method, originally proposed by Ciccotti *et al* (57, 58), combines simulations in equilibrium and nonequilibrium conditions (Figure S5). It allows for the direct computing of the evolution of the dynamical response of a system to an external perturbation. The rationale for the D-NEMD can be described as follows: if an external perturbation is applied to an equilibrium simulation and, by doing so, a parallel nonequilibrium simulation is started, then the response of the protein to the perturbation can be straightforwardly extracted using the Kubo-Onsager relation (for more details, see (44, 46)).

The perturbation used here, the instantaneous removal of LA from all three FA sites, is the same as in previous D-NEMD simulations of non-glycosylated spikes (28–30). This perturbation takes the system out of equilibrium, and creates a driving force for changes to occur as the protein responds to the perturbation and relaxes back towards equilibrium. LA removal from the FA sites triggers the structural response of the protein as it adapts to an empty FA site. Analysis of the evolution of the structural changes reveals the mechanical and dynamical coupling between the structural elements involved in response to LA removal and identifies the allosteric pathways connecting the FA site to the rest of the protein. The evolution of the structural response of the protein is extracted using the Kubo-Onsager relation (44, 45, 57, 59) from the difference between the equilibrium and nonequilibrium trajectories at equivalent points in time (Figure S7). The response obtained for each pair of equilibrium and nonequilibrium simulations is averaged over all 210 trajectories here, hence reducing noise (44, 45), and allowing the statistical significance of the responses to be assessed from the standard error of the mean (Figure S7-S9) (44).

LA removal initiates a complex chain of structural changes that are, over time, propagated within the protein. The deletion of the LA molecules immediately triggers a structural change in the FA site, which contracts as the sidechains of the residues lining it move closer to each other, filling the space once occupied by the LA molecule (Figure S10). The changes in the FA site are then swiftly transmitted to well-defined regions of the protein, notably the NTD, RBM and FP-surrounding regions (Figures 2 and S11-S12).

**Figure 2.**
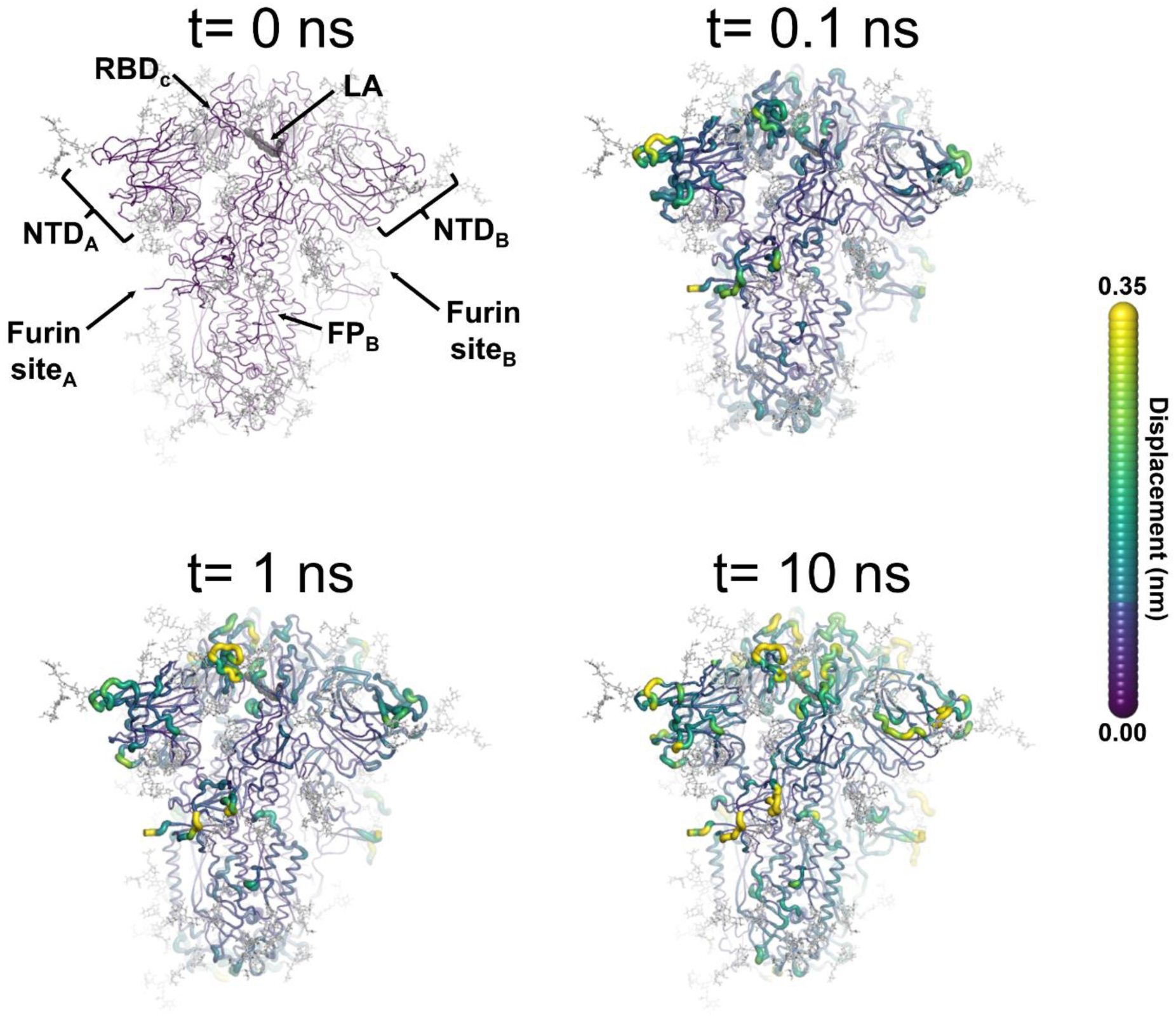
Structural response of the glycosylated spike to LA removal. The average Cα displacements 0.1, 1 and 10 ns after LA removal from the FA binding sites are shown, mapped onto the starting structure for the equilibrium simulations. The norm of the average Cα displacement vector between the D-NEMD apo and equilibrium LA-bound simulations was calculated for each residue using the Kubo-Onsager relation (44, 45, 57, 59). The final displacement values are the averages obtained over the 210 pairs of simulations (Figures S7-S9). The cartoon thickness and structure colours (scale on the right) indicate the average Cα-positional displacement. Each RBD, NTD, furin site and FP are subscripted with their chain ID (A, B or C). Glycans are shown as light grey sticks, whereas the dark grey spheres highlight the position of the LA molecule. The FA site shown in this figure is FA site 1, which is located at the interface between chains C and A (see Figures S11-S12 for the protein responses from the viewpoint of FA site 2 and 3, respectively). The responses of all three FA sites are qualitatively similar, with the same motifs and sequence of events observed.

The cascade of events observed here for the fully glycosylated ancestral spike mirrors that of the non-glycosylated protein (28–30), with LA removal triggering immediate structural changes in the FA site, which are then transmitted to key regions of the protein, including to the FP-surrounding region which is more than >40 Å away from the FA site. Figure S13 shows a strong positive correlation between the responses obtained for the non-glycosylated and glycosylated spikes, underscoring the overall similarity between our current and previous D-NEMD simulations. The largest differences in the responses between the glycosylated and non-glycosylated spikes are found around the furin recognition site (Figure S13). This is unsurprising as the furin site is cleaved in the current (glycosylated) simulations but remains uncleaved in the previous non-glycosylated simulations. The furin cleavage/recognition site is a polybasic four-residue insertion located on a solvent-exposed loop at the S1/S2 junction (16, 17). This site is important for the activation of the spike (60), and its presence affects viral infectivity (e.g. (29, 60–63)).

The evolution of the response of the spike to LA removal reveals the pathways through which structural changes propagate from the FA site to functional motifs (e.g. motifs involved in membrane fusion and antigenic epitopes (Figures 3 and S14-S15)). These pathways lie mainly within the protein and are generally similar to those previously found for the non-glycosylated spike (28–30). The structural changes induced by LA removal, which start in the FA site (mainly in the P337-A348 and S366-A372 regions of one monomer and T415-K417 of the other one), are rapidly transmitted to the rest of the RBD. The R454-K458 region is particularly important as it mediates the transmission of the structural changes to the A475-C488 segment in the RBM (Figure 3A).

**Figure 3.**
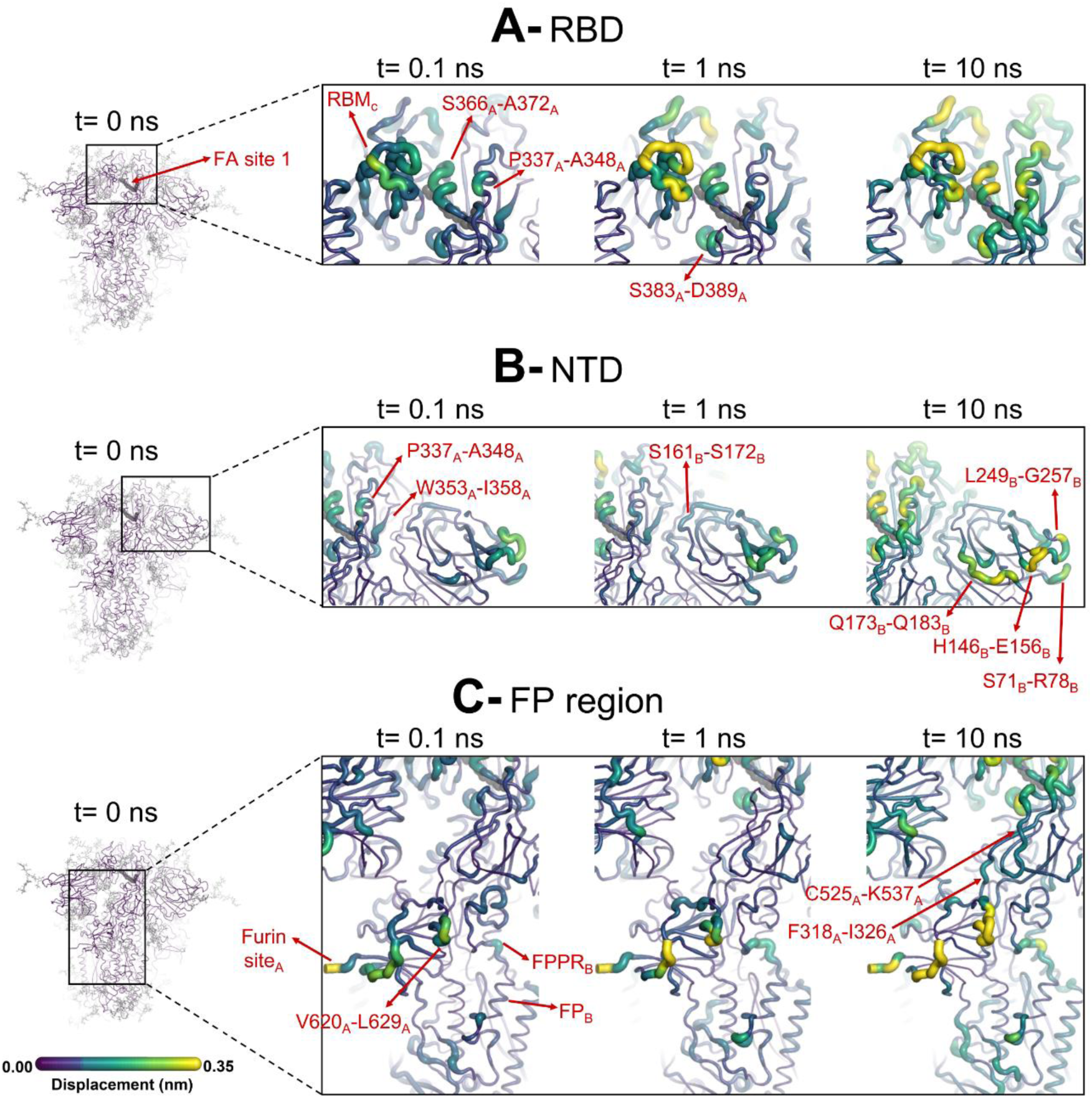
Structural responses of functional regions of the spike. Close-up view of the structural response of the RBD (**A**), NTD (**B**) and FP surrounding regions (**C**) to LA removal. The FA site shown here is FA site 1, located at the interface between chains C and A (see Supporting Figures S14-S15 for the responses of the other two FA sites, which are similar). Structure colours and cartoon thickness indicate the average Cα displacement values. Each region is subscripted with its chain ID (A, B or C). The dark grey spheres show the FA binding site. In the images representing the spike at *t=*0 ns (left side images **A**, **B** and **C**), the glycans are shown as light grey sticks. Glycans were omitted from the figures showing the responses at *t=*0.1, 1 and 10 ns to facilitate visualization. For more detail, see the legend of Figure 2.

In addition to the amplitude of the structural changes induced by LA removal, the average directions of the motion can also be computed by determining the average displacement vector of Cα atoms (54) between the equilibrium and nonequilibrium trajectories at equivalent time points (Figures 4 and S17-S20). Indeed, the amplitude of the structural changes in Figures 2-3, S11-S12 and S14-S15, corresponds to the norm of the average Cα displacement vector. Upon LA removal, the spike regions that form the FA site show responses with well-defined directions. These segments include P337-A348 and S366-A372, which are regions with residues whose side chains form the FA pocket (Figures 4 and S17-S20). Soon after LA deletion, P337-A348 and S366-A372 (P337_A_-A348_A_ and S366_A_-A372_A_ in FA site 1, P337_B_-A348_B_ and S366_B_-A372_B_ in FA site 2 and P337_C_-A348_C_ and S366_C_-A372_C_ in FA site 3) move inwards towards the FA site. These motions collectively reflect the contraction of the FA site upon LA removal. The directions of RBM motions are more diverse, with two of the RBMs, namely RBM_C_ (Figure S18) and RBM_B_ (Figure S19), displaying an upward movement and the third showing an opposite downward one (RBM_A_) at *t=*10 ns (Figures 4 and S20).

**Figure 4.**
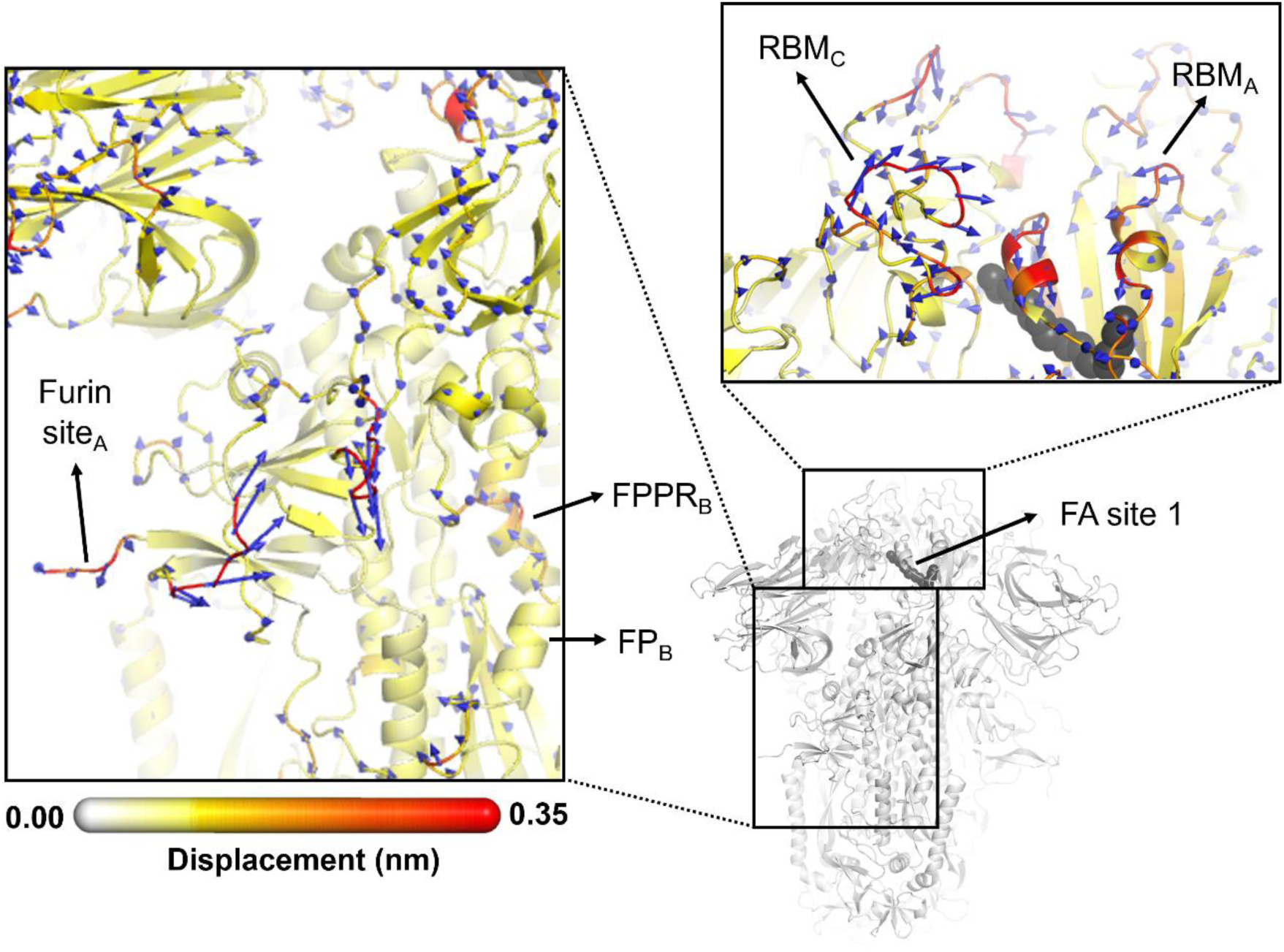
Directions of the structural responses of the RBD and FP-surrounding regions to LA removal. The average Cα displacement vectors at *t=*10 ns are shown. These vectors were determined by averaging Cα displacement vectors between the equilibrium and nonequilibrium trajectories over the 210 replicas. Vectors with a length ≥0.1 nm are displayed as blue arrows with a scale-up factor of 10. The average displacement magnitudes are represented on a white-yellow-orange-red scale. The dark grey spheres represent the FA site. This figure shows the directions of the responses around FA site 1, which is located at the interface between chains C and A (see Supporting Figures S19-S20 for the direction of the motions around the other two FA sites).

The ancestral SARS-CoV-2 spike has two glycans located on the RBD: *N-*glycans at positions N331 and N343 (35). The N343 glycan is particularly interesting because it is situated immediately after one of the regions that transmits structural changes from the FA site, namely P337-F342 (Figure 3). As well as being close to the FA site, this glycan can also directly bridge the two neighbouring RBDs (Figure S16), which may strengthen the connection between the FA site and these regions. N343 has been shown to play a role in the RBD opening mechanism by acting as a gate for the change from the ‘down’ to the ‘up’ conformation (32).

The structural changes induced by LA removal are also swiftly propagated to the NTD via P337-F342, W353-I358, and C161-P172 (Figures 3 and S14-S15). Despite variations in amplitude, the structural response to LA removal is consistent across the three FA sites, exhibiting the same motifs and sequence of events for signal propagation to the NTDs. The response of the protein, which starts in the P337-A348 segment in the FA site, is transmitted to W353-I358, C161-P172 and then to several antigenic epitopes located in the periphery of the NTDs (64). The external regions showing high displacements, namely S71-R78, H146-E156, Q173-Q183 and L249-G257, are all part of an antigenic supersite in the distal-loop region of the NTD (65). In particular, the GTNGTKR motif in S71-R78, besides being an antigenic epitope, has also been suggested to be involved in binding to other receptors, such as sugar receptors (10).

The conformational response of the Q173-Q183 is of particular interest as this region forms a pocket that binds heme and the tetrapyrrole products of its metabolism (66, 67). The structural response initiated in the FA site propagates through the NTD, reaching the Q173-Q183 segment (Figures 3, 5 and S14-S15). This region, which is located in the distal face of the NTD, forms the entrance of an allosteric site (Figure 5) that has been shown to bind heme (66) and its metabolites biliverdin and bilirubin (67). X-ray crystallography, cryo-EM and mutagenesis experiments together with modelling show that heme and biliverdin bind to a deep cleft in the NTD (66, 67) gated by the Q173-Q183 flexible loop (Figure 5B). Physiological concentrations of biliverdin suppress binding of some neutralizing antibodies to the spike (67). Such data suggested a new mode of immune evasion of the spike via the allosteric effect of biliverdin/heme binding (67). In our D-NEMD simulations, two of the three Q173-Q183 regions (in chains A and B) show well-defined outward motions in response to LA removal (Figures 5 and S20). Our results show a clear connection between the FA and biliverdin/heme allosteric sites via internal conformational motions despite the two sites being >30 Å apart. This connection, captured by the D-NEMD approach, is remarkable and illustrates the complexity of the potential allosteric modulation of the spike. These results suggest that the presence of heme or its metabolite in the NTD site affects the internal networks and how dynamic and structural changes are transmitted to and from the FA site. The presence of heme/biliverdin may modulate the response of the spike to fatty acids and vice-versa, potentially affecting the rate and/or affinity of binding of the molecules to their respective allosteric sites. This apparent connection between allosteric sites is worthy of further investigation.

**Figure 5.**
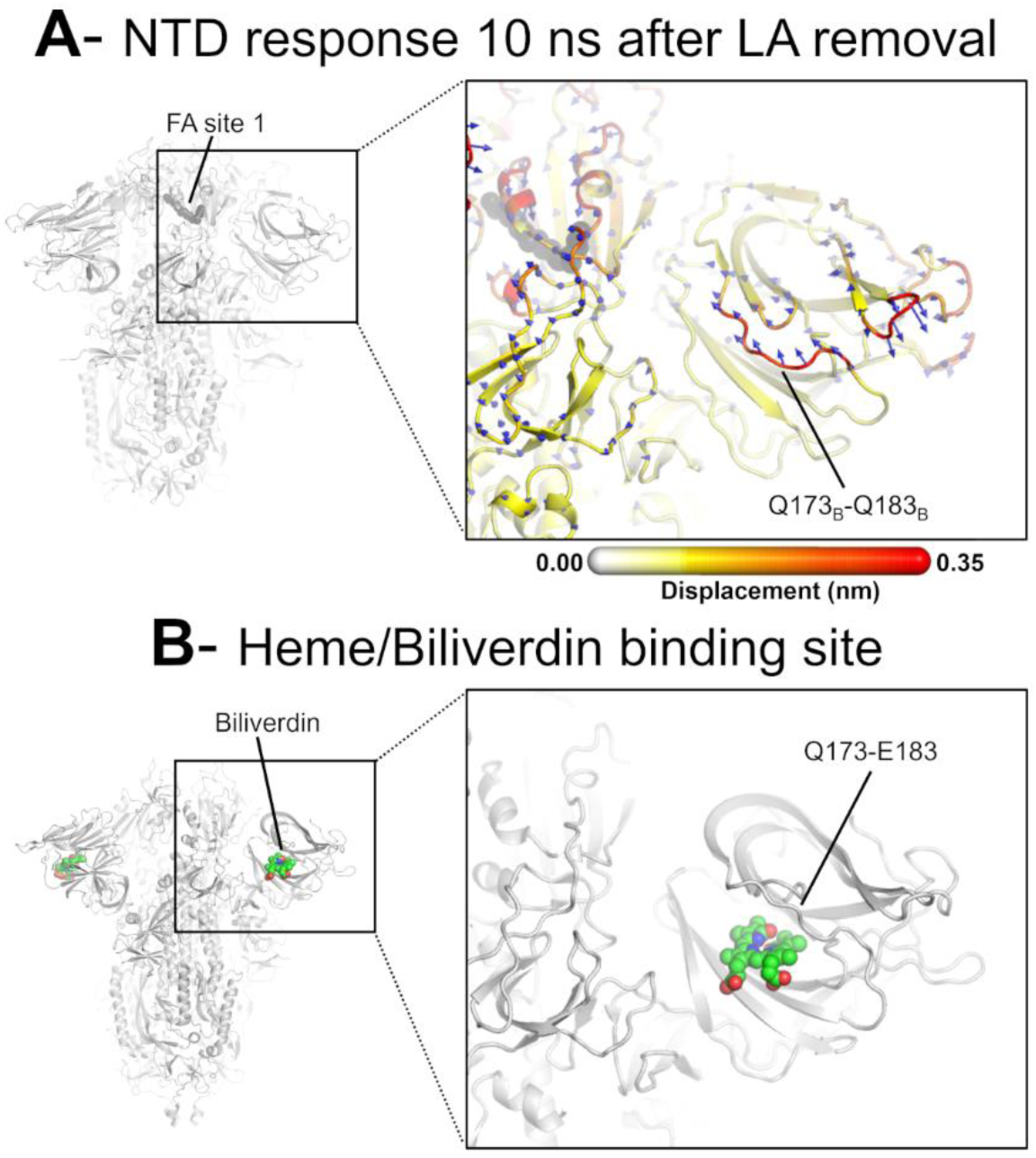
D-NEMD displacement vectors show a connection between the FA site and the heme/biliverdin binding site in the NTD. (**A**) View of the NTD 10 ns after LA removal, focusing on the heme/biliverdin binding site (66, 67) (which is not occupied in the simulations here). Note that the Q173-Q183 segment, which contains residues forming the heme/biliverdin binding site, shows an outward motion upon LA removal. The magnitudes of the displacements are represented on a white-yellow-orange-red colour scale. Vectors with a length ≥0.1 nm are displayed as blue arrows with a scale-up factor of 10. The dark grey spheres represent the FA site. This figure shows the direction of the structural responses around FA site 1 (see Figure S21 for the direction of the motions in the other two FA sites). (**B)** Cryo-EM structure showing the heme/biliverdin binding site in the NTD (PDB code: 7NT9)(67). The protein is coloured in grey. The biliverdin molecules are shown with spheres.

There are eight *N-*glycans on the NTD, linked to N17, N61, N74, N122, N149, N165, N234 and N282 (35). Of these, four (N74, N122, N149 and N165) are located in or close to the segments that respond to FA site occupancy (Figure S16). N74, N122 and N149 are involved in shielding the spike protein from the host immune system (33), whereas the role of N165 and N234 goes beyond shielding: they are involved in the stabilization of the RBD ‘up’ conformation (33).

The structural changes induced by LA removal are not restricted to the RBD and NTD: they are also transmitted to several regions far from the FA site, notably the furin site at the S1/S2 boundary, the S2’ protease recognition and cleavage site, R815, located the S2 subunit (immediately upstream the fusion peptide), V620-L629 loop and FP proximal region, FPPR (Figures 3 and S14-S15). The structural responses starting in the FA site quickly propagate downwards to the furin cleavage sites and V620-L629 loop via the F318-I326 and C525-K537 segments (Figures 4 and S18-S20). The furin cleavage site, which harbours a polybasic motif containing multiple arginine residues, is located at the boundary between the S1 and S2 subunits and more than 40 Å from the FA site. The furin cleavage site is important for protein activation, and its removal reduces viral infectivity (29, 60, 62, 68). The addition of extra positively charged residues near the furin cleavage site, as observed in several variants, has been suggested to increase proteolytic processing (69), and has been shown to increase the rate of binding and the affinity of glycosaminoglycans such as heparin and heparan sulfate for this area (70). In our previous work, the comparison of the D-NEMD responses to LA removal between variants of concern has also suggested that the addition of extra positive flanking charges, which is observed in some variants (such as P681R in Delta and N679K in Omicron), strengthens the allosteric connection between the FA and furin cleavage site (28). Overall, the allosteric effects observed here for the glycosylated ancestral spike are qualitatively similar to those in the non-glycosylated protein: the same regions are connected to the FA site and are affected significantly by ligand removal from this site (28–30). This indicates that the glycans on the exterior of the protein do not substantially affect the internal allosteric communication pathways within the spike.

The structural changes starting in the FA site are transmitted to the furin cleavage site and V620-L629, and from there, over time, propagated to the FPPR and S2’ cleavage site (Figures 4 and S18-S20). In addition to being an epitope for neutralizing antibodies (71, 72), the S2’ cleavage site is also crucial for infection (63, 73). This proteolytic site is located immediately before the hydrophobic FP in the S2 subunit, and its cleavage is mediated by the transmembrane protease serine 2 (TMPRSS2) after binding to the host receptor (73). The FPPR is located after the FP, and it is thought to have a functional role in membrane fusion by mediating the transitions between pre- and post-fusion structures of the protein (18). Upon removal of LA, the residues of the FPPR in direct contact with the FP show a well-defined response upwards in two of the chains, namely chains B and C, and an outwards motion in chain A (Figures 4 and S19-S20). FPPR_B_ and FPPR_C_, located closer to FA sites 1 and 2 respectively, show an upward movement towards the C-terminal domain 1 (CTD1), whereas FPPR_A,_ which is nearer to FA site 3, exhibits a lateral outward motion. The CTD1 has been suggested (based on cryo-EM structures) to be a structural relay between RBD and FP, sensing displacement on either side (18)). Interestingly, the chains displaying FPPR motion towards CTD1, notably chains B and C, also exhibit an RBM upward movement away from the body of the spike. The direction of the S2’ motion observed in the D-NEMD simulations is diverse, with two of the sites (S2’_A_ and S2’_B_) displaying a motion towards the FP and one showing an opposite movement away from the FP (S2’_C_) after t = 10 ns (Figures 4 and S17-S20).

The spike contains several complex *N*- and *O*-glycans in or close to the furin and S2’ cleavage sites, FPPR, F318-I326, C525-K537 and V620-L629 (35). All three monomers contain one *O*- and two *N*-glycans (at positions T323, N616 and N657) close to the pathway that connects that FA site to the furin cleavage site and FP regions (Figure S16). Monomer A also contains an additional *O*-glycan linked to S323 (Figure S16). Notably, interactions between the *O*-glycans S323 and T325 and the *N*-glycan at N234 create a direct connection between the NTD and the F318-I326 region of the same monomer (Figure S16). This glycan ‘link’ may facilitate and enhance the transmission of structural changes within an individual subunit. The glycan at position N234 has also been suggested to play a mechanical role in the spike infection mechanism by helping to stabilize the RBD in the ‘up’ conformation (33).

Cross-correlation analysis was performed for the equilibrium and D-NEMD simulations (similarly to references (28, 55)) to identify the coupled regions in the protein, including those with motions connected to the FA site. In Figure S22, the dark and light blue regions represent high and moderate negative correlations between the Cα atoms in the protein, and red and orange regions correspond to high and moderate positive correlations. Negative correlation values indicate that the atoms are moving in opposite directions, whereas atoms systematically moving along the same direction show strong positive correlations. Overall, the cross-correlation maps computed from the equilibrium and D-NEMD trajectories show similar patterns, with the former exhibiting a subtle increase in the correlations between the FA sites and RBDs (Figure S22). This increase indicates that binding a fatty acid molecule, such as LA, to the FA site reinforces the connection between this site and other parts of the protein.

The cross-correlation maps also show positive correlations between each FA site and two of the three RBDs in the protein. This is because each FA site sits at the interfaces between every two neighbouring RBDs (e.g. FA site 1 is formed by residues from subunits A and C). Low to moderate negative and small positive correlated motions are observed between the FA site and the NTDs and fusion peptide regions, respectively (Figures S22). To visualize these motions, the statistical correlations for R408 and K417 (two FA site residues able to form salt-bridge interactions with the carboxylate head group of LA) were mapped on the protein structure (Figure S23). Figure S23 shows the patterns of movement described above and the regions whose motions are connected to the FA site. Interestingly, some segments forming the signal propagation pathways, such as R454-K458 in all three monomers and C525-K537 in monomers B and C can also be identified from the cross-correlation analysis, showing moderate to high correlations with the FA site (Figures S22-S23).

To assess whether the substitutions, deletions and insertions seen in various SARS-CoV-2 variants lie on the allosteric communication pathways identified, we overlapped, in the same 3D structure, the sequence variations (using spheres to highlight the position of the changes) with the D-NEMD responses reported above (Figure S25). In Figure S25, changes within the allosteric pathways are indicated by red spheres, while those within 0.6 nm of any atom (i.e., both main and sidechain atoms) forming the paths are highlighted in dark blue. The results are interesting: 22 of the 77 amino acid positions per chain that differ in the Alpha, Beta, Gamma, Delta and Omicron (BA.1, BA.2, BA.4, BA.5, BQ.1.1 and XBB.1.5) variants directly map onto the allosteric communication pathway identified using D-NEMD. A further 28 out of the 77 variations are in direct contact with, or very close proximity to, these networks. H655Y (present in Gamma and all Omicron sub-variants), T547K (in Omicron BA.1), D614G (in Alpha, Betta, Gamma, Delta and all Omicron sub-variants), W856K (in Omicron BA.1) and S982A (in Alpha) are all examples of mutations close to the communication pathways, which may influence the connection to the FA site. These mutations are responsible for the previously observed differences in allosteric behaviour between SARS-CoV-2 variants (28).

Sequence alignment of the original spike and the Alpha, Beta, Gamma, Delta and Omicron (BA.1, BA.2, BA.4, BA.5, BQ.1.1 and XBB.1.5) variants (Figure S24) shows that several of the mutations, deletions and insertions present in the variants are located either in or near the pathways identified here. Furthermore, some variants, such as Beta, Gamma, and Omicron, contain residue substitutions at the FA site. For example, the lysine in position 417 in the original spike is mutated to asparagine in Beta and Omicron and threonine in the Gamma variant. Another example is arginine 408 in the Ancestral protein, which has been replaced by asparagine in several Omicron sub-variants. As future variants emerge, it will be of interest to establish whether mutations lie in the FA and heme/biliverdin (66, 67) sites or along the allosteric pathways described here. Differences in allosteric behaviour and regulation in the spike are likely to be of functional relevance and may be useful in understanding differences between SARS-CoV-2 variants. Our previous work using D-NEMD revealed significant differences in the allosteric responses of spike variants to LA removal (28). The substitutions, insertions, and deletions in the variants affected both the amplitude of the structural responses and the rates at which these rearrangements propagate within the protein (28). The allosteric connections identified in Alpha were generally similar to the ancestral protein, whereas Delta exhibited increased connections between the FA site and the furin cleavage site but diminished links to V622–L629 (28). Omicron displayed significant changes in the NTD, RBM, and furin cleavage site, with stronger couplings observed between the FA site and these regions (28).

## Conclusions

D-NEMD simulations show important allosteric effects in the fully glycosylated ancestral SARS-CoV-2 spike. These simulations identify the pathways that link the FA site with functional regions (regions of the spike involved in membrane fusion, antibody recognition and allosteric modulation). The D-NEMD simulations show the structural responses resulting from LA removal and demonstrate connections between the FA site and: the RBM; an antigenic supersite in the NTD; the allosteric heme/biliverdin binding site (66, 67); the furin site; and, the FP-surrounding region (including S2’ cleavage site and, the FPPR) (Figure S26). The connection between the FA site and the RBM is mediated by residues R454-K458, while the transmission of structural changes from the FA site to the NTD involves residues P337-F342, W353-I358, and C161-P172. The allosteric pathways connecting the FA site to the furin site and FP-surrounding region involve the segments containing residues F318-I326 and C525-K537. Notably, more than 65% of the substitutions, deletions and additions in the Alpha, Beta, Gamma, Delta, and Omicron variants are located either in or close to the allosteric pathways identified using D-NEMD.

Furthermore, our D-NEMD results reveal an unexpected connection between the FA site and a second allosteric site known to bind heme and its metabolites (66, 67). It will be of interest to understand how heme/biliverdin binding affects the dynamics and structural changes of the spike, and links to the FA site, and potentially other allosteric sites (74). While the effects of the apparent coupling between the heme/biliverdin site and the FA site remain to be investigated, this work has reinforced the ability of the D-NEMD approach to find allosteric sites and to map communication pathways between sites (44, 46). The results here further point to the complex allosteric effects in the SARS-CoV-2 spike, of potential functional relevance.

Comparison with previous D-NEMD simulations of the non-glycosylated spike (28–30) shows that the presence of glycans on the exterior of the protein, does not qualitatively change the cascade of events connecting the FA site to the rest of the spike. Some glycans influence the allosteric pathways, facilitating the transmission of the structural changes within and between subunits. For example, the interactions between the glycans linked to N234, T373 and S375 can create a direct connection between the NTD and the F318-I326 region of the same monomer, thus helping the propagation of structural changes within the monomer. These results shed new light on the roles of glycans and further emphasise their potential in modulating the functional dynamics of the spike.

## Acknowledgements

ASFO was supported at the University of Bristol by a Fellowship from Oracle for Research and is a BBSRC Discovery Fellow ([BB/X009831/1]). This work is part of a project that has received funding from the European Research Council under the European Horizon 2020 research and innovation programme (PREDACTED Advanced Grant Agreement no. 101021207) to AJM. IB and CS are investigators of the Wellcome Trust (202904/Z/16/Z, 206181/Z/17/Z, 221708/Z/20/Z). ADD is a member of the G2P2-UK National Virology Consortium funded by the Medical Research Council (MRC)/UKRI (Grant MR/Y004205/1). AJM and ASFO thank the Biotechnology and Biological Sciences Research Council (BBSRC) grant number BB/W003449/1). All MD simulations were carried out using the Oracle Public Cloud Infrastructure (https://cloud.oracle.com/en_US/iaas) under an award to AJM and ASFO from Oracle for Research for COVID-19 research. We thank EPSRC for providing ARCHER/ARCHER2 time through a COVID-19 rapid response call via HECBioSim (hecbiosim.ac.uk). Data analysis was conducted using the facilities of the Advanced Computing Research Centre at the University of Bristol (https://www.bris.ac.uk/acrc/). We also thank the Bristol UNCOVER Group and the University of Bristol for support. This work was also supported in part by NSF RAPID MCB-2032054, an award from the RCSA Research Corp., a UC San Diego Moore’s Cancer Center 2020 SARS-COV-2 seed grant to REA.

## Data availability statement

All D-NEMD simulation data (including input and trajectories files) will be openly available from the MolSSI/BioExcel COVID-19 public data repository for biomolecular simulations of COVID proteins (https://covid.molssi.org/simulations/).

## Competing interests

The authors declare competing interests: CS and IB report shareholding in Halo Therapeutics Ltd related to this Correspondence.

## Notes

### Competing Interest Statement

The authors declare competing interests. Christiane Schaffitzel and Imre Berger report shareholding in Halo Therapeutics Ltd related to this Correspondence.

### Summary of Updates

- Added several new analyses to the manuscript (cross-correlation analysis and sequence analysis) - Several new figures were added to SM to support the new analyses performed - the list of authors was updated

