## Supporting Information for "Allosteric modulation by the fatty acid site in the glycosylated SARS-CoV-2 spike"

#### Materials and methods

##### Model for the glycosylated spike with linoleic acid bound

A model of the fully glycosylated ectodomain of the ancestral (also known as wild type, 'early 2020', or original) spike with three closed RBDs and one linoleate (LA) molecule bound in each free fatty acid (FA) binding site (i.e. three linoleate molecules) was created based on the cryo-EM structure 7JJI (1) following protocols applied and tested previously for the ancestral spike without linoleate, with the same glycosylation profile as in the work by Casalino *et al.* (2). The model for the glycosylated spike-LA complex contains 15 disulphide bonds per trimer and is cleaved at the furin protease cleavage site located at S1/S2 interface. Similar models for the fully glycosylated closed spike have been widely tested and used in a wide range of applications (e.g. (2-5)).

The ancestral spike model here contains 22 N- and two O-glycosylation sites per monomer, as in previous work. Note, however, that these sites have been found to be heterogeneously populated in different experimental studies (e.g. (6, 7)). The spike model used as starting point for this work reflects this heterogeneity, with asymmetric site-specific glycosylation profiles derived from the glycoanalytic data reported by Watanable *et al.* for the N-glycans (6) and Shajahan *et al.* for the O-glycans (7). This means that glycan occupancy and composition differ between the three monomers. A detailed description of the glycans is available in ref. (2).

##### Equilibrium simulations

All equilibrium MD simulations were performed using the CHARMM36m all-atom force field (8, 9). The simulation conditions and protocols were the same as those applied successfully previously in refs. (2, 3). Starting with a closed spike head conformation in which glycan N74 was fully outstretched, the spike-LA complex was placed in a rectangular box (19.5nm x 21.5nm x 20.5nm, ensuring at least 1 nm separation from the x and y edges of the box, and 1.5nm separation from the z edges of the box) solvated with TIP3P water and with 150 mM NaCl. Special care was taken to solvate with a sufficiently large water box to avoid self-interaction energies by glycans crossing periodic boundaries.

For the solvated spike-LA complex, we conducted the following minimisation, heating, and equilibration protocols in triplicate with NAMD2.14 (10) on AmaroLab local machines: parameters for linoleate in these early preparatory simulations were taken from Paramchem.org (11-14), consistent with CGenFF parameters, and passed in to NAMD with a stream file. All following steps were performed for each replicate. The waters and ions (774,333 water atoms, 701 Na atoms, 687 Cl atoms) were minimised for 10,080 steps using the default conjugate gradient energy minimisation algorithm in NAMD, during this time protein and glycan atoms were held fixed with Lagrangian constraints. From minimised coordinates, water and ion atoms were then progressively heated in the NVT ensemble from 10 K to 310 K over the course of 120.96 ps, wherein temperature was increased by 25 K every 10.08 ps (timestep 1.0 fs/step). Once temperature reached 310 K, an additional 766.08 ps of equilibration simulation (timestep 1.0 fs/step).

Following water and ion minimisation and heating, we released all Lagrangian constraints, added positional restraints on protein and glycan atoms (force constant 1 kcal/mol/Å^2) and performed a quick conjugate gradient minimisation of the whole restrained system for 2520 steps. We then randomly reinitialised velocities for all atoms at 310 K and performed 252,000 steps of NpT equilibration (timestep=2.0 fs/step) with a Nosé-Hoover Langevin piston driven barostat (pressure = 1.0325 bar, piston temperature = 310 K, useFlexibileCell = yes, useGroupPressure = yes). Following this restrained NpT relaxation, we conducted 50 ns (timestep 2fs/step) of unrestrained (all positional restraints removed) NpT equilibration, with fixed box dimensions (useFlexibleCell = no). From the final frame of the 50ns NpT equilibrations, we used CHARMM-GUI (15-17) to convert NAMD psf and coordinate files to GROMACS compatible itp and gro files. Three replicate simulations, each 750 ns, were performed using GROMACS (18) using the CHARMM36m all-atom force field (8, 9). The simulation conditions were the same as in (2). All GROMACS unrestrained equilibrium MD simulations were performed on Oracle Cloud Infrastructure (OCI) using compute nodes consisting of 8×NVIDIA A100 tensor core GPUs, and 64 AMD Rome CPU cores. Using 64 CPU cores and 8 GPUs per simulation allowed us to achieve a performance of ~70 ns/day.

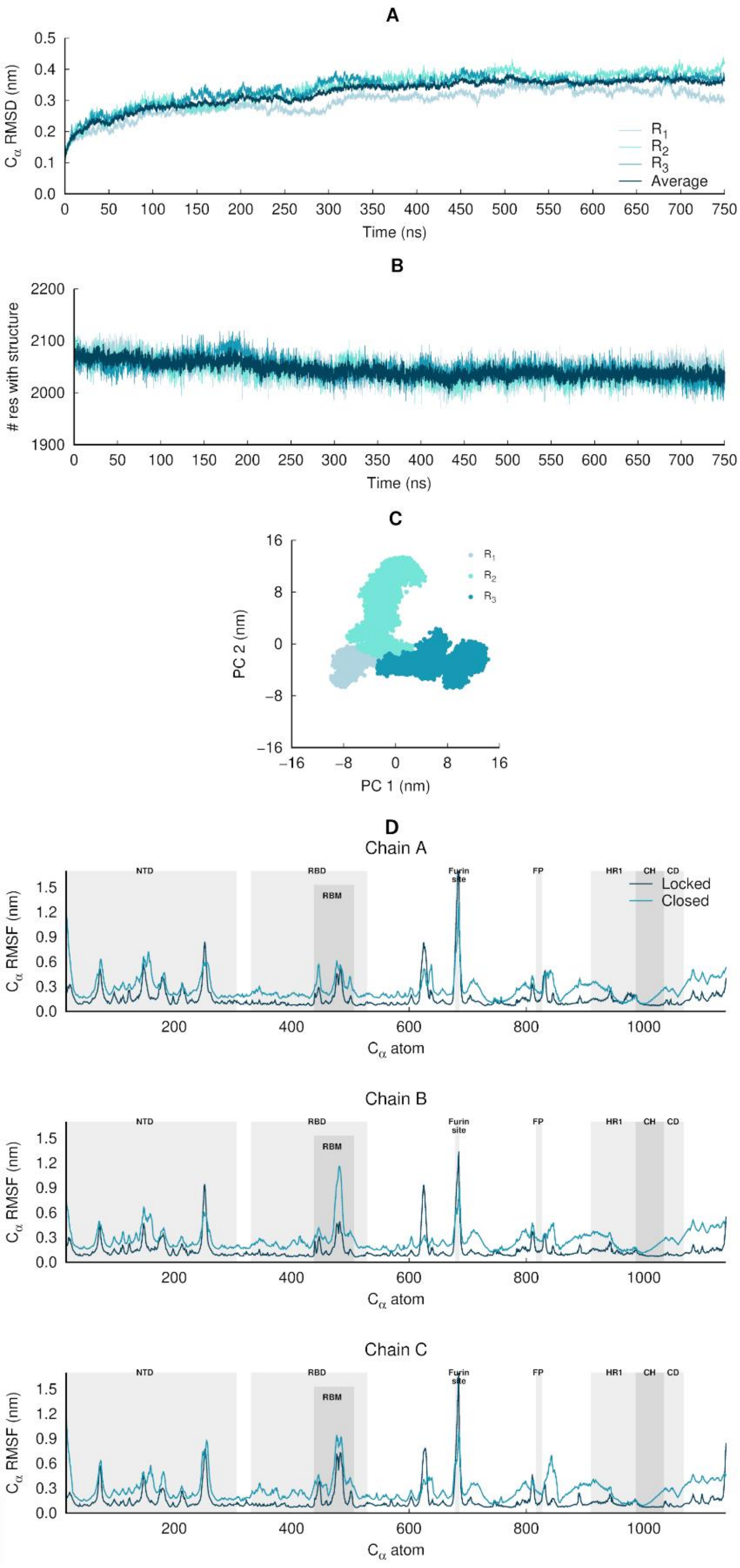


**Figure S1. Structural stability, equilibration and sampling of the equilibrium simulations of the glycosylated SARS-CoV-2 ancestral spike**. (**A**) Temporal evolution of the $C_\alpha$ root mean square deviations (RMSD) relative to the starting structure. The dark blue line corresponds to the average $C_\alpha$ RMSD, averaged over all three replicate simulations (namely $R_1$, $R_2$ and $R_3$). These results indicate that the systems are stable over the simulation time. (**B**) Time evolution of the spike's secondary structure content (assigned by DSSP (19)), showing numbers of residues assigned to α-helix, $3_{10}$-helix, 5-helix, β-sheet and β-bridge secondary structure classes. The light blue lines correspond to the time evolution of the number of residues with secondary structure in the individual replicate simulations, whereas the dark blue line corresponds to the average number of residues over all three replicates. (**C)** Principal component analysis (PCA) of the three replicates, for all $C_\alpha$ atoms. All replicates were combined for the PCA, with one conformation per 100 ps per replicate (in a total of 22501 frames). (**D**) Average $C_\alpha$ root mean square fluctuations (RMSF) for the glycosylated ancestral spike in the locked (with LA bound) and closed (without LA) states. The $C_\alpha$ RMSF was calculated using the entire equilibrium trajectories and averaged across all replicates for each state (three replicates for locked spike and three for the closed one). The trajectories for the closed glycosylated spike were taken from Casalino *et al.* (2). The grey boxes identify key regions in the protein, namely the N-terminal domain (NTD), receptor-binding domain (RBD), receptor-binding motif (RBM), fusion peptide (FP), heptad repeat 1 (HR1), central helix (CH), connector domain (CD). Please note that the fusion-peptide proximal region (FPPR) is situated immediately before the FP. Please zoom in on the image for detailed visualisation.

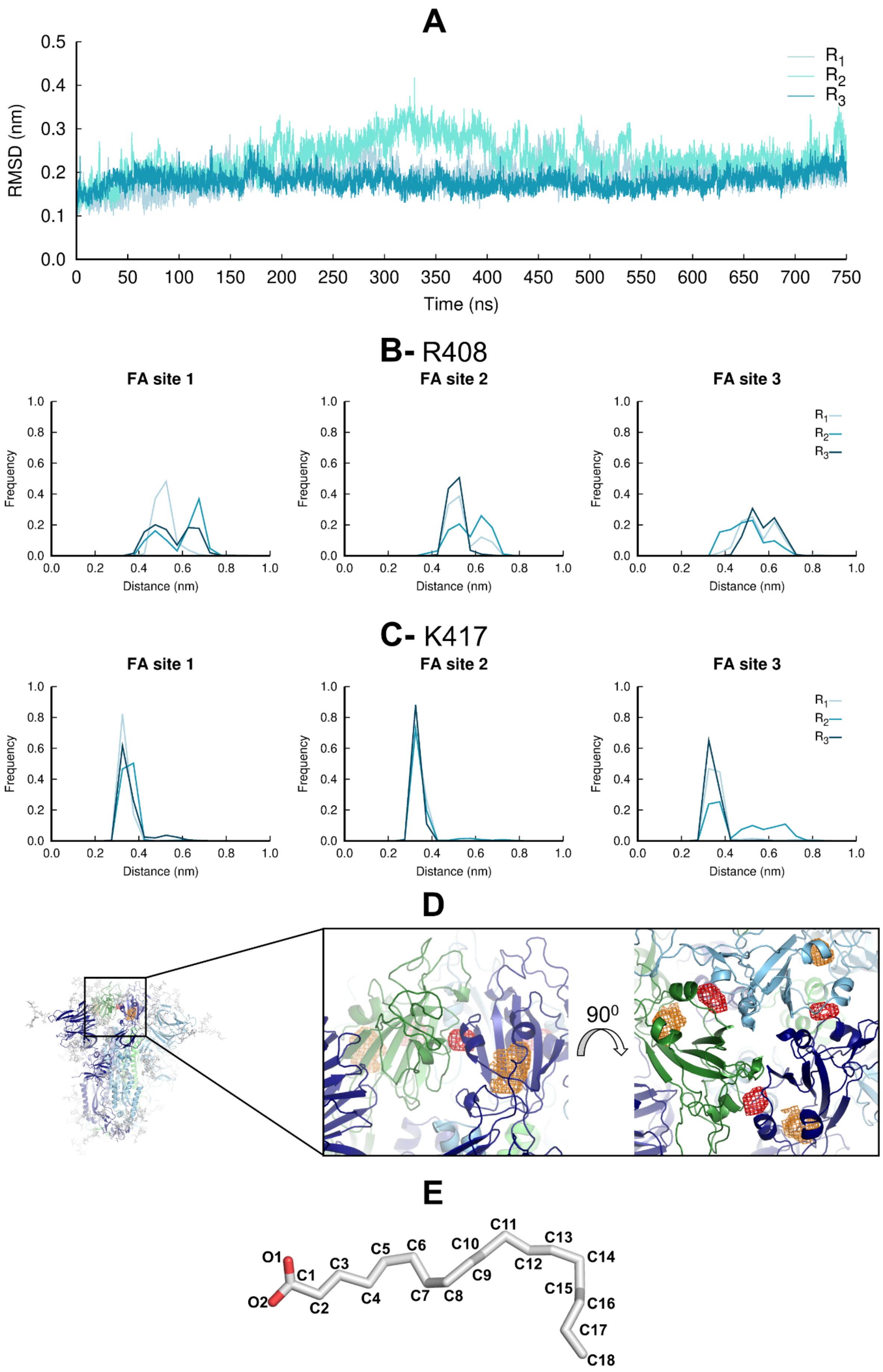
A
RMSD (nm)
Time (ns)
R1
R2
R3
B- R408
C- K417
FA site 1
FA site 2
FA site 3
Frequency
Distance (nm)
D
90⁰
E
O1
O2
C1
C2
C3
C4
C5
C6
C7
C8
C9
C10
C11
C12
C13
C14
C15
C16
C17
C18

**Figure S2. Motions and interactions of the LA molecules during the equilibrium MD simulations**. (**A)** Average RMSD of the LA molecules for each replica relative to the starting model. The RMSD values are the average over the three LA molecules. No LA molecule exits the FA site during the equilibrium MD simulations, consistent with high-affinity binding at this site (20, 21). **(B** and **C)** Distributions of the distances between the negatively charged carboxylate group of LA and the positively charged side chain of R408 (**B**) and K417 (**C**) in each FA site. (**D)** Probability density maps (with a 0.00001 Å$^{-3}$ contour) for the carboxylate carbon C1 (red mesh) and the aliphatic carbon C18 (orange mesh). See panel E for FA structure and atom label. The probability density maps were determined using 7501 frames per replicate. The structure used as the starting point for the simulations is shown with cartoon. Each monomer in the spike homotrimer is shown in a different colour: dark blue, light blue and green. **(E)** Structure of LA, with labels in all nonhydrogen atoms.

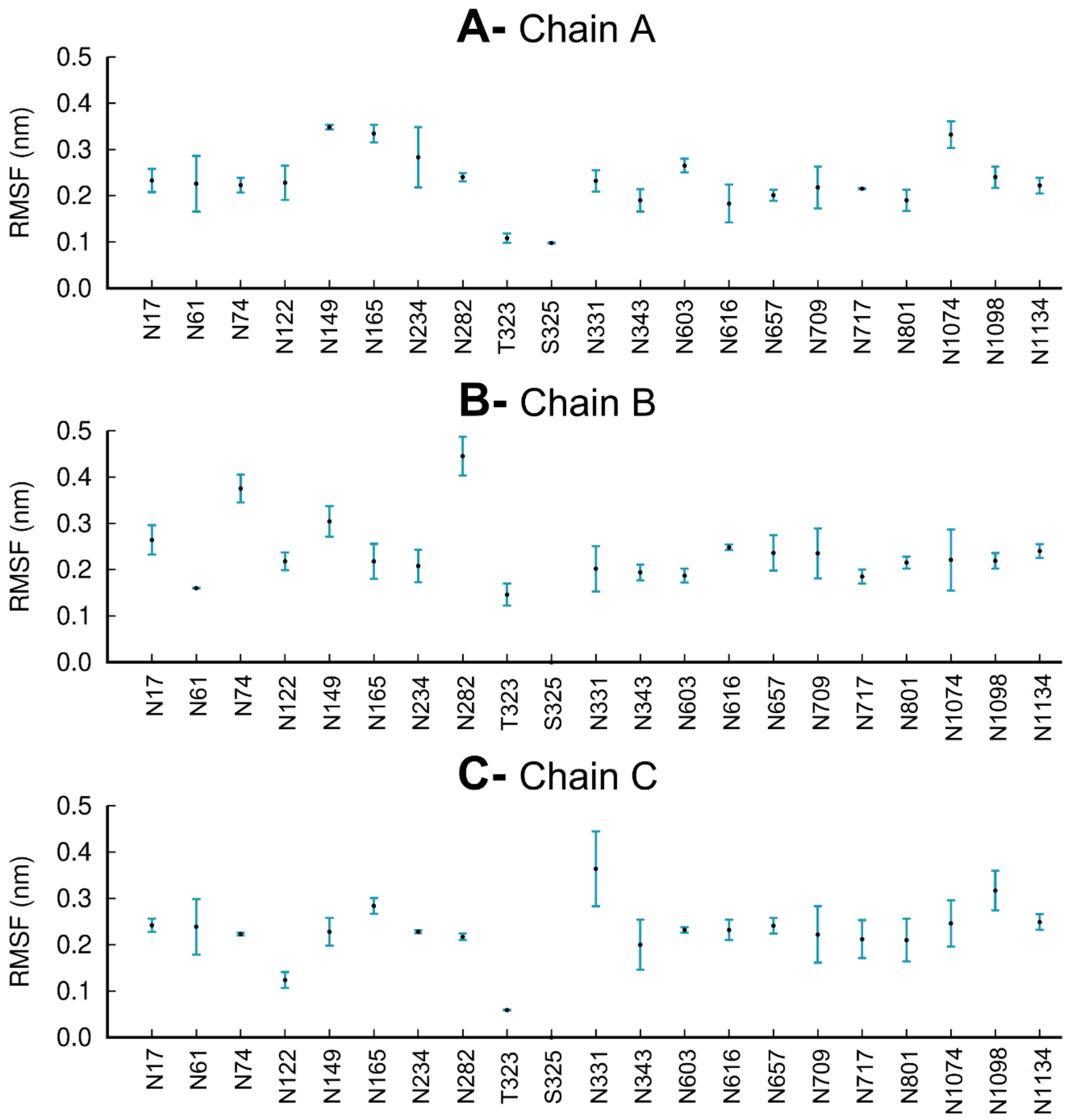


**Figure S3. Average RMSF of each glycan for chains A (A), B (B), and C (C) over the three replicate equilibrium MD simulations**. The vertical blue lines represent one standard deviation. Note that glycosylation site S325 is only occupied in chain A, with no glycans present in this site in chains B and C, similarly to Casalino *et al.* (2).

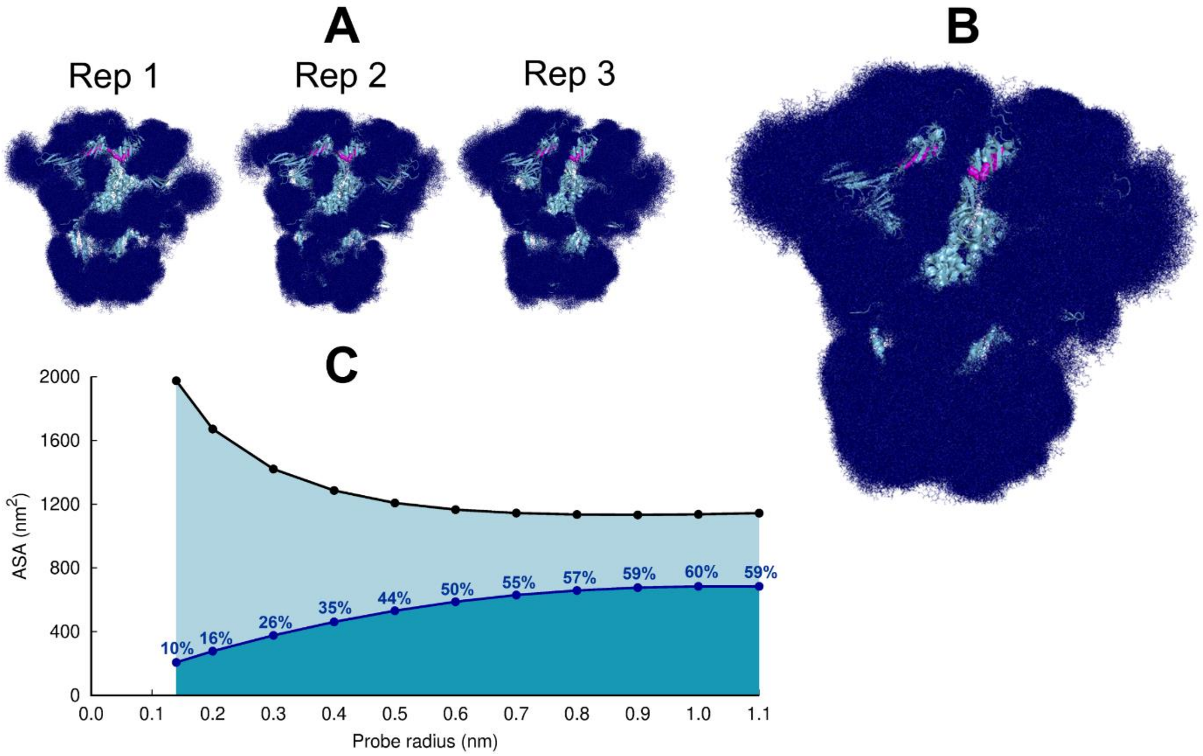


**Figure S4. Glycan shielding of the spike**. (**A**) Overlapping of the conformations adopted by the glycans during the simulation for each individual replica. The position of the glycans in 376 frames (one frame every 2 ns) are shown with dark blue sticks. (**B**) Overlapping of the glycan conformations in all three replicas (in a total of 1128 frames). The protein is shown as a light blue cartoon whereas the glycans are the dark blue sticks. The magenta spheres represent the LA molecules. (**C**) Solvent accessible surface area of the protein and the area shielded by glycans at multiple probe radii. The probe radius ranges from 0.14 nm (corresponding to a water molecule) to 1.1 nm (corresponding to a small antibody molecule). The values are averaged across all replicas. The area shielded by the glycans corresponds to the dark blue line, whereas the black line represents the accessible surface area of the protein without glycans (similarly to e.g. ref (2, 22)).

#### Dynamical nonequilibrium molecular dynamics (D-NEMD) simulations

We apply a dynamical approach to nonequilibrium molecular dynamics (D-NEMD) simulations that was proposed by Ciccotti *et al.* more than forty years ago (23, 24). This approach combines MD simulations in equilibrium and nonequilibrium conditions and allows computation of the evolution of the dynamic response of a system to an external perturbation (25-27).

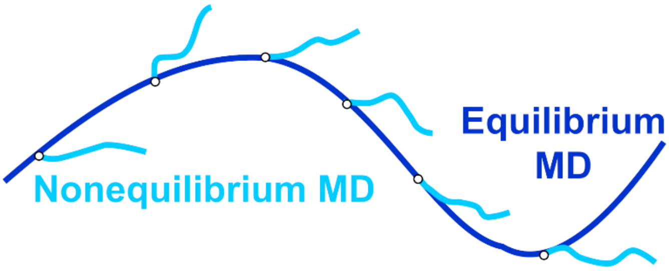

**Figure S5- Schematic representation of the D-NEMD approach.** Equilibrium MD trajectories (dark blue line) provide the initial distribution of conformations for the nonequilibrium simulations at time t=0 (light blue lines).

The rationale for the D-NEMD approach can be described as follows: if an external perturbation (e.g. removal of a ligand) is added to a simulation sampling an equilibrium state and, by doing so, a parallel nonequilibrium simulation is started (Figure S5), then the structural response of the protein to the perturbation can be measured by comparing the equilibrium and nonequilibrium trajectories at equivalent points in time by using the Kubo-Onsager relation (as long as enough sapling is gathered (25-27)). This approach has the advantage that the statistical significance of the response can be easily assessed, and the associated errors calculated and made as small as desirable by increasing the number of nonequilibrium trajectories. Determining the statistical errors associated with the responses (through, e.g., the determination of the standard error of the mean) is essential to test if the sampling gathered is sufficient (25-27). Here, the standard error of the mean was calculated for each average Cα displacement value at times 0.1, 1 and 10 ns after the removal of LA (see Figures S7-S9).

Generally, multiple (tens to hundreds) D-NEMD simulations are needed to achieve statistically significant results for biomolecular systems (for examples, see (25, 26)). The length of the D-NEMD simulations performed (usually 5-10 ns long) reflects a balance between the computational resources available and the number of replicates needed to achieve statistically significant responses.

Here, a large set of D-NEMD simulations was performed to study the structural response of the fully glycosylated, cleaved (cleaved at the furin recognition site) ancestral spike to LA removal. 210 nonequilibrium simulations (70 simulations per replicate), each 10 ns long, were carried out. The procedure used to set up and analyse the nonequilibrium simulations is illustrated in Figure S6. The 210 starting configurations for the D-NEMD simulations were obtained from the equilibrated part of the equilibrium LA-bound trajectories (Figure S6). Conformations were taken every ten ns to begin the nonequilibrium simulation: for each, all of the LA molecules bound to the three FA sites were (instantaneously) removed. The resulting nonequilibrium apo system was then simulated for 10 ns (Figure S6). The simulation conditions for the nonequilibrium simulations were the same as LA-bound equilibrium simulations described above. The perturbation used here, namely the removal of LA from the FA sites, is the same as in previous work (28-30). This perturbation is designed to force the system out of equilibrium, thus creating a driving force and forcing structural changes to propagate within the protein (26).

The response of the spike to LA removal from the FA sites was computed using the Kubo-Onsager relation (23, 26, 27, 31), by calculating the displacement of each $C_\alpha$ atom between the equilibrium and nonequilibrium simulations at equivalent points in time (Figure S6). For each time point, the $C_\alpha$ atom displacement vector was averaged over the 210 replicas (Figures S7-S9). The resulting average vector indicates the average direction of the response of the residues upon the perturbation; the norm of the average displacement vector gives the amplitude of the responses. The statistical significance of the responses was assessed by determining the standard error of the mean (Figures S7-S9).

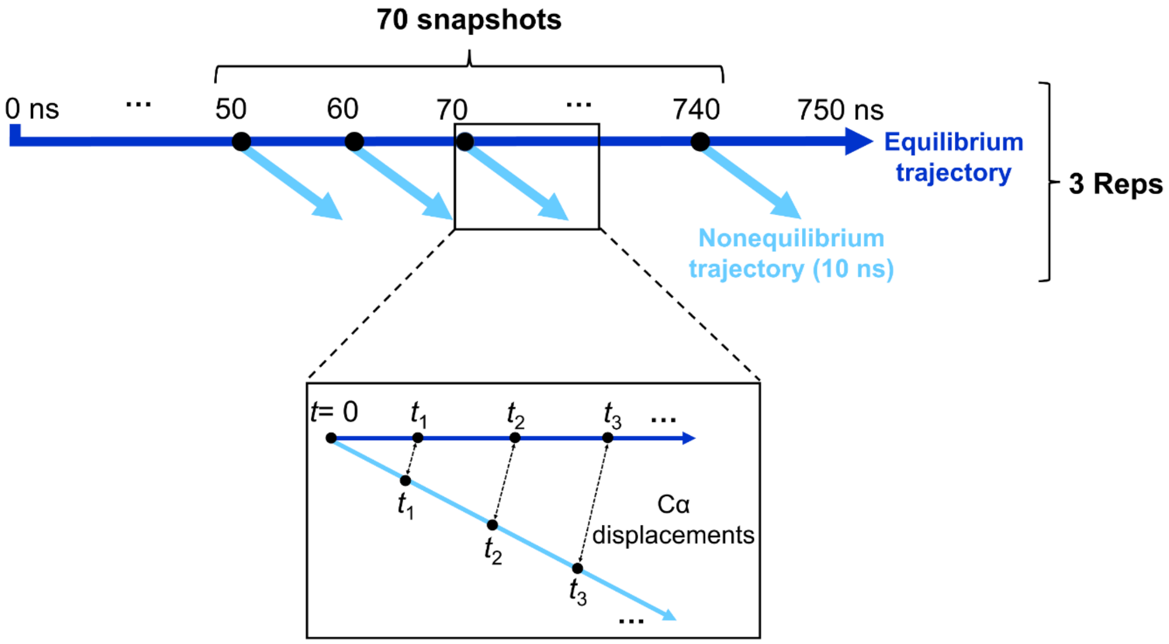


**Figure S6. Schematic of the procedure used to set up and analyse the D-NEMD simulations here**. Three equilibrium MD simulations, 750 ns each, were performed for the fully glycosylated, cleaved (with cleavage at the S1/S2 interface) spike in the closed state. These equilibrium trajectories were then used to generate starting structures for the short apo nonequilibrium simulations. From the equilibrated part of each LA-bound simulation (from 50-750 ns), conformations were extracted every ten nanoseconds, and the perturbation was introduced. Each D-NEMD simulation was run for 10 ns. The Kubo-Onsager (23, 26, 27, 31) relation was used to extract the response of the system to LA annihilation from the FA pockets: for each pair of equilibrium LA-bound and D-NEMD apo trajectories, the displacement of each $C_\alpha$ at equivalent times (namely 0, 0.1, 1 and 10 ns) was determined and averaged over all 210 pairs of simulations.

### Supporting Results

Figures S7-S9 show the D-NEMD responses of the protein's $C_{\alpha}$ atoms at different time points.

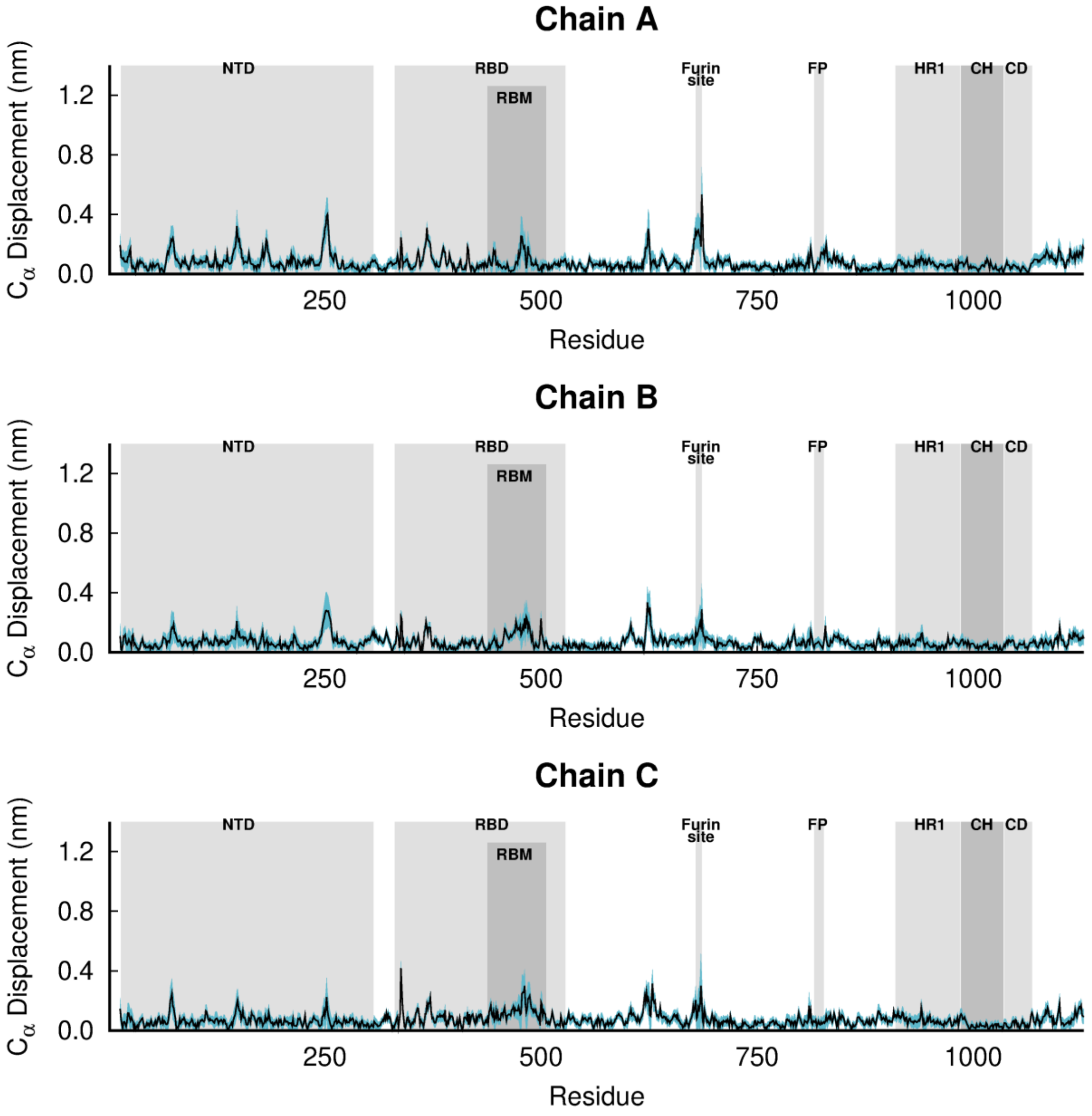


**Figure S7. D-NEMD average $C_{\alpha}$-positional displacement and corresponding standard errors 0.1 ns after LA removal from the FA sites**. The average deviations were determined using the Kubo-Onsager relation (23, 26, 27, 31) for the pairwise comparison between the nonequilibrium apo and equilibrium LA-bound simulations. The averages were calculated over the 210 pairs of simulations. The light blue shaded region represents the standard error of the mean. The positions of some key structural motifs are highlighted in grey, namely the N-terminal domain (NTD), receptor-binding domain (RBD), receptor-binding motif (RBM), fusion peptide (FP), heptad repeat 1 (HR1), central helix (CH), connector domain (CD). Please note that the fusion-peptide proximal region (FPPR) is situated immediately before the FP. Please zoom in on the image for detailed visualisation.

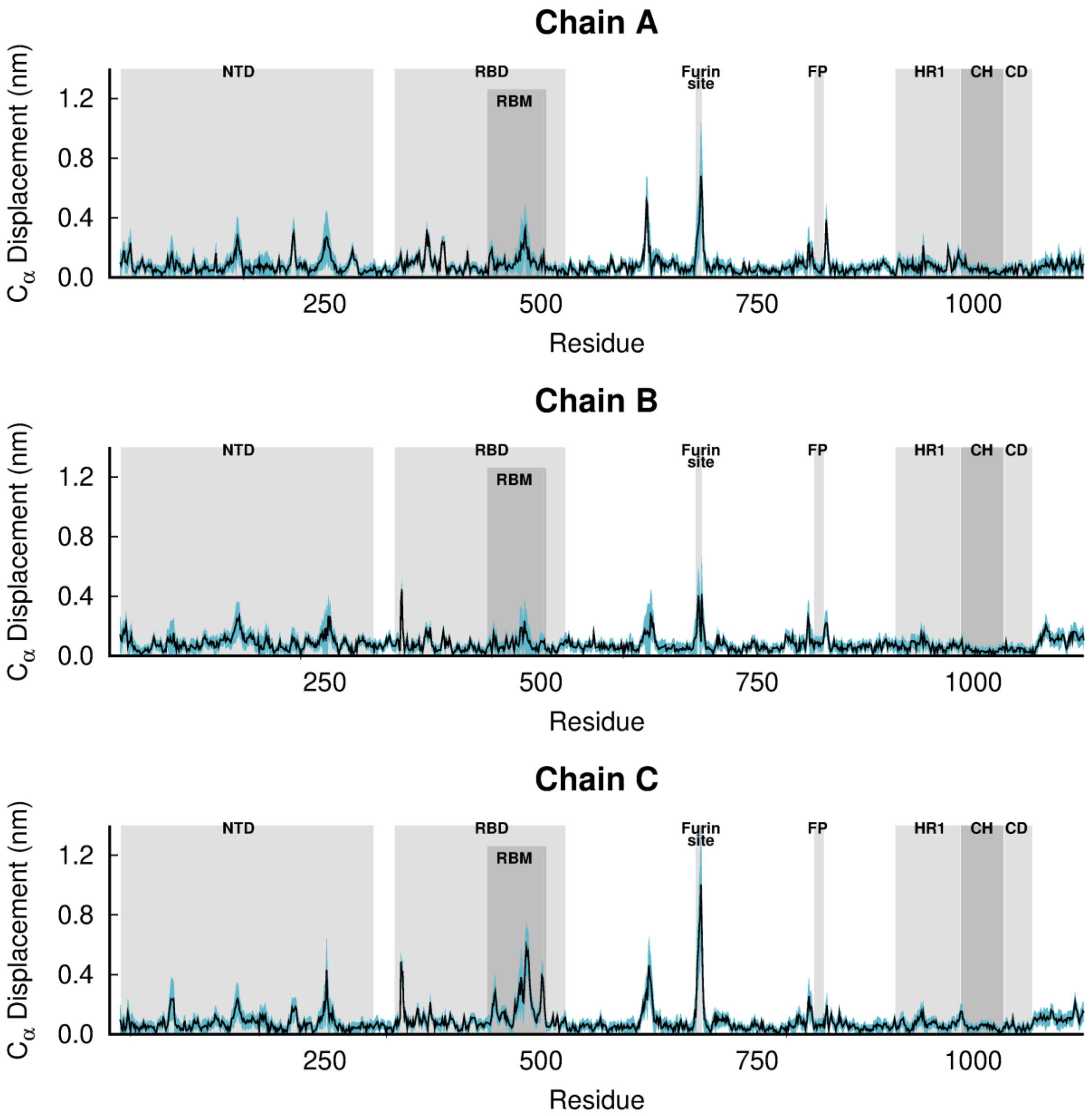


**Figure S8. D-NEMD average $C_\alpha$-positional displacement and corresponding standard errors 1 ns after LA removal from the FA sites**. For more details, see the legend of Figure S7.

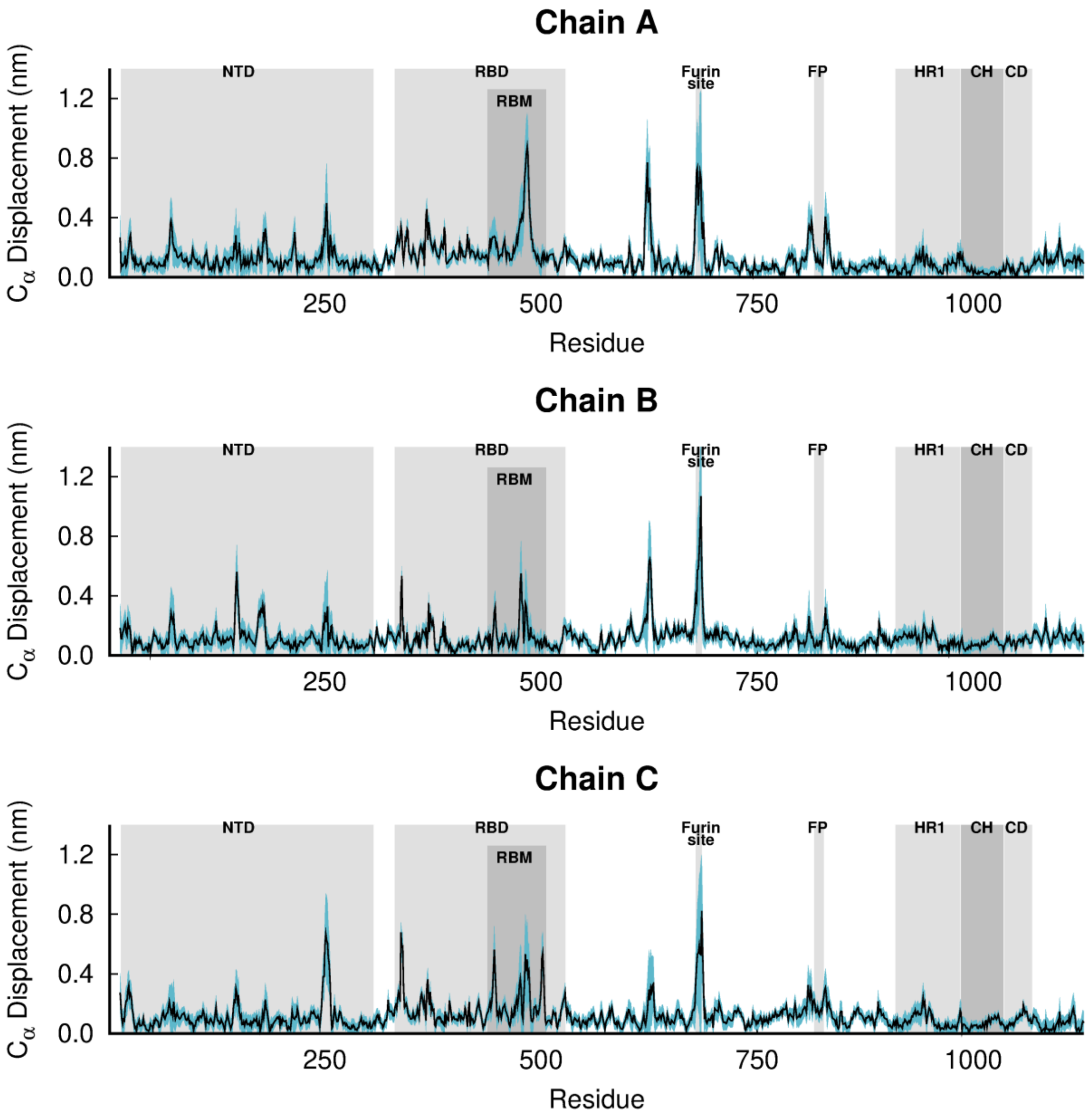


**Figure S9. D-NEMD average $C_\alpha$-positional displacement and corresponding standard errors 10 ns after LA removal from the FA sites**. For more details, see the legend of Figure S7.

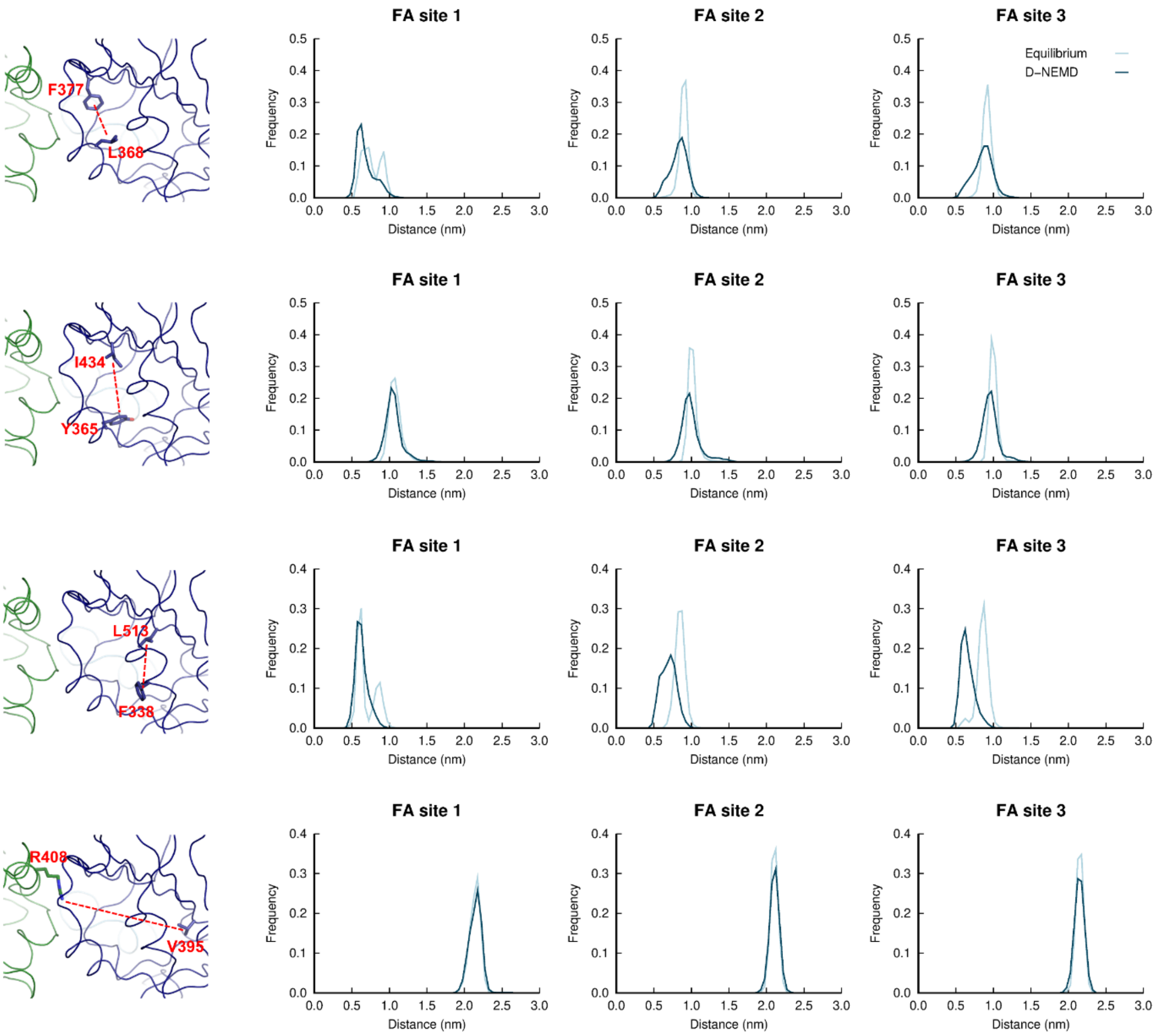


**Figure S10. Distributions of distances between: L368-F377; Y365-I434; F338-L513; and, R408-V395 in the equilibrium LA-bound and D-NEMD apo simulations**. Histogram of the distance between the centre of mass of the sidechains of: L368 and F377; Y365 and I434; F338 and L513; and, R408 and V395 in the equilibrium and D-NEMD simulations. The equilibrium histogram (light blue line) displays distance data from the three equilibrium trajectories from 0-750 ns. The D-NEMD histogram (dark blue line) contains the distance data from all 210 D-NEMD trajectories. The images of the FA site shown on the left show example conformations of the residues shown in the histograms on the right.

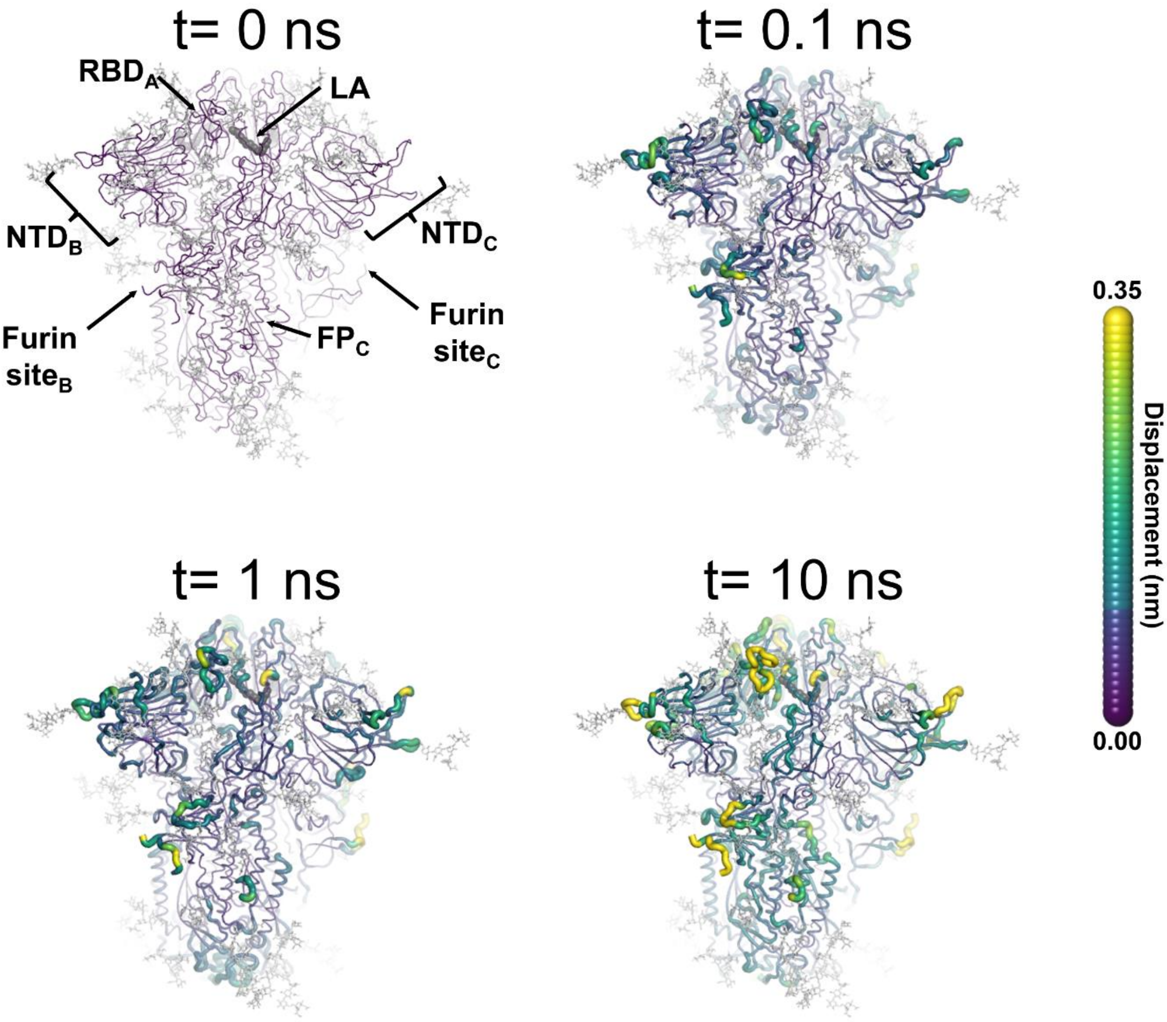


**Figure S11. Structural response of spike to the removal of LA**. This figure shows the protein's response from the viewpoint of FA site 2, which is located at the interface between chains A and B. Chain ID (A, B or C) is indicated in subscript for each RBD, NTD, furin site and FP). Mapping of the average $C_\alpha$-positional displacement after LA removal from the FA binding sites. The norm of the average $C_\alpha$ displacement vector between the D-NEMD apo and equilibrium LA-bound simulations was calculated for each residue. The final displacement values correspond to the average obtained over the 210 pairs of simulations. The $C_\alpha$ average displacements at *t*=0, 0.1, 1 and 10 ns are mapped onto the starting structure for the equilibrium simulations. Structure colours indicate the average $C_\alpha$-positional displacement (according to the scale on the right). Glycans are shown as light grey sticks, and the dark grey spheres highlight the FA binding site. The displacements in this figure can be compared with those shown in the main text Figure 2 (showing the changes from the FA site 1 side).

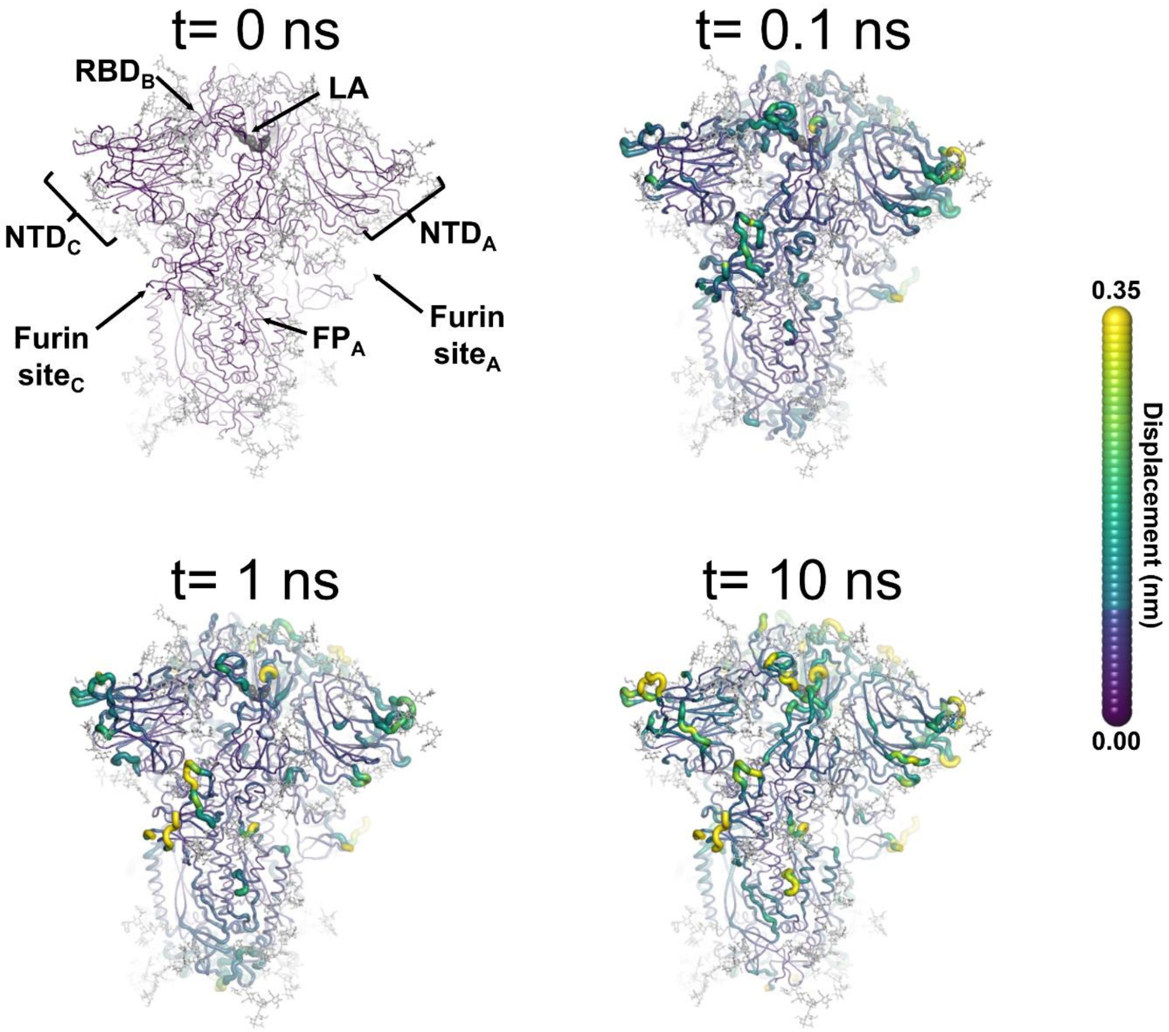


**Figure S12. Structural response of spike to the removal of LA**. This figure shows the protein's response from the viewpoint of FA site 3, which is located at the interface between chains B and C. Chain ID (A, B or C) is indicated in subscript for each RBD, NTD, furin site and FP. Mapping of the average $C_\alpha$-positional displacement after LA removal from the FA binding sites. The norm of the average $C_\alpha$ displacement vector between the D-NEMD apo and equilibrium LA-bound simulations was calculated for each residue. The final displacement values correspond to the average obtained over the 210 pairs of simulations. The $C_\alpha$ average displacements at *t*=0, 0.1, 1 and 10 ns are mapped onto the starting structures for the equilibrium simulations. Structure colours indicate the average $C_\alpha$-positional displacement (according to the scale on the right). Glycans are shown as light grey sticks, and dark grey spheres highlight the FA binding site. The displacements in this figure can be compared with those shown in the main text Figure 2 (showing the changes from the FA site 1 side).

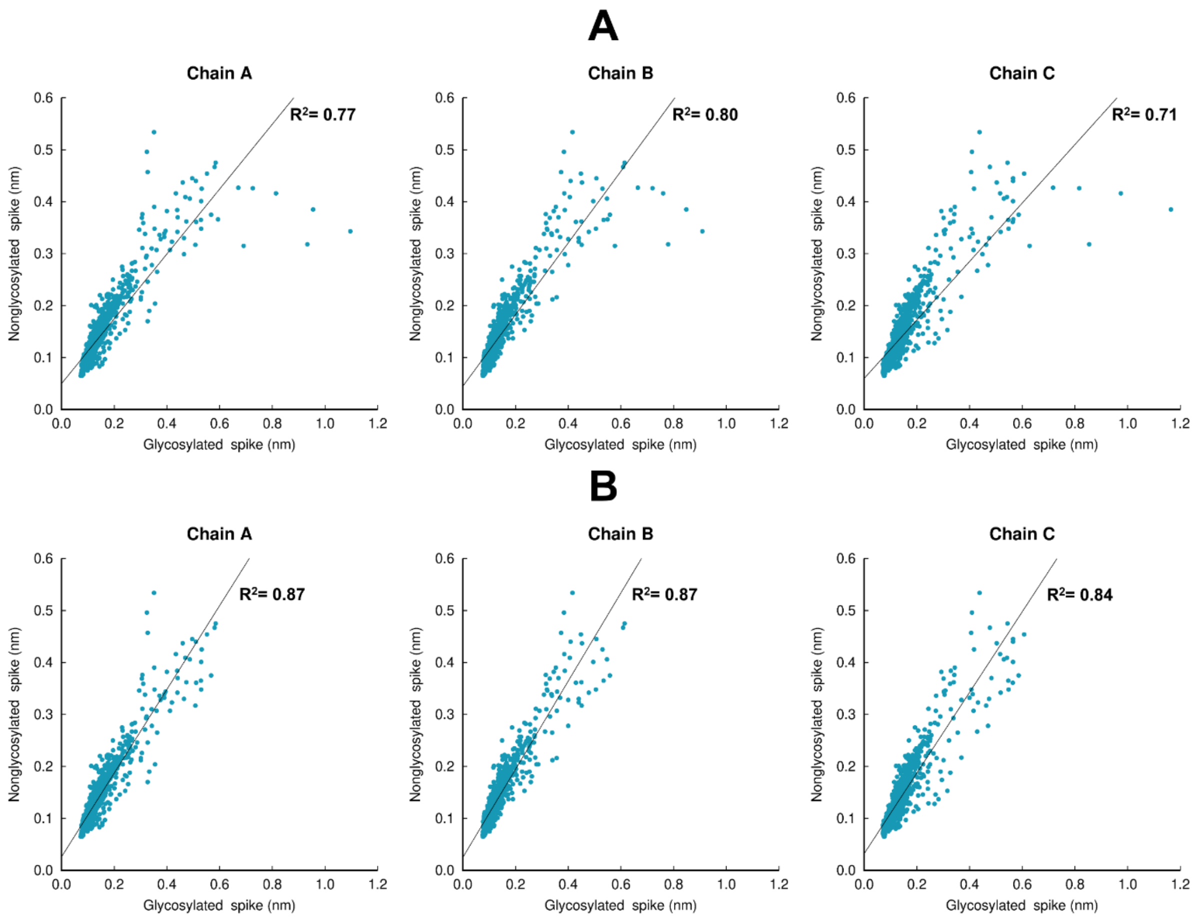


**Figure S13. Structural response of the glycosylated spike compared to the non-glycosylated protein**. (**A**) Scatter plots comparing the responses for the complete head region of the spike (residues 13-1140). (**B**) Scatter plots comparing the protein response excluding the residues around the furin recognition site (residues 680-690). Panels A and B show the values for the average $C_\alpha$-positional deviations 10 ns after LA removal from the FA sites. The average deviations for the non-glycosylated ancestral spike are taken from our previous work (28). To properly compare the responses, the average $C_\alpha$-positional deviations for the glycosylated spike was calculated using the same approach as in (28). The coefficient of determination value for the linear regression for each chain is shown. The comparison between the coefficient values for panels A and B shows that the largest differences in the response between the glycosylated and non-glycosylated spikes are located in the furin recognition site at the S1/S2 interface, which is cleaved in the glycosylated protein here and uncleaved in the non-glycosylated spike in previous simulations (28); this difference should be noted in comparing the responses here.

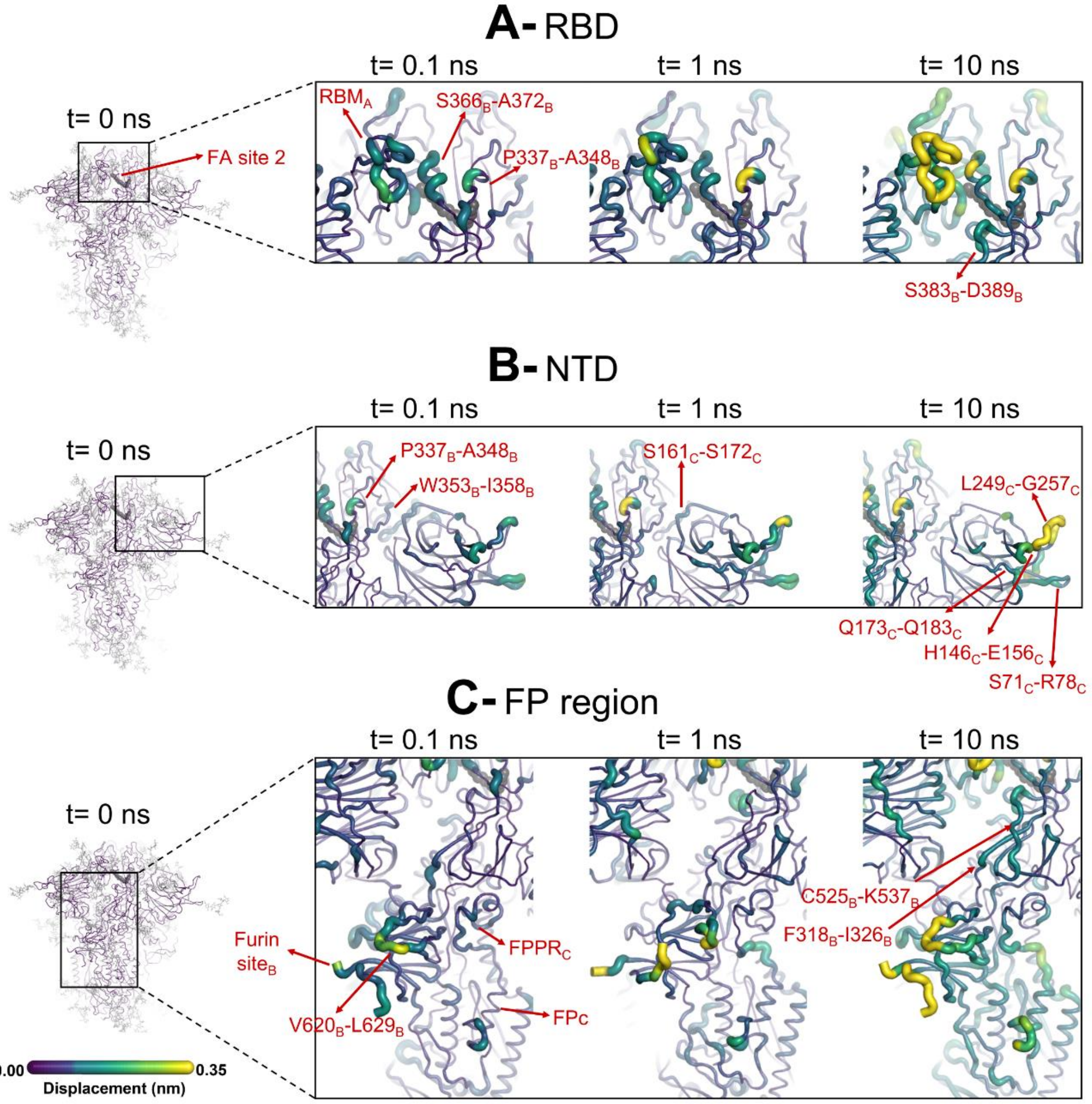


**Figure S14. Evolution of the D-NEMD structural response from the viewpoint of FA site 2** (located at the interface between chains A and B) to LA removal. Close-up view of the structural response of the $RBD_{AB}$ (**A**), $NTD_{C}$ (**B**) and $FP_{C}$ surrounding regions (**C**) to LA removal. Chain ID (A, B or C) is indicated in subscript. The average $C_{\alpha}$ displacements at *t*=0, 0.1, 1 and 10ns after LA removal are mapped onto the starting structure for the equilibrium simulations. The structure colours, and cartoon thickness, indicate the average $C_{\alpha}$-displacement values. The dark grey spheres highlight the FA binding site. The glycans are omitted from the panels showing the responses at *t*=0.1, 1 and 10 ns.

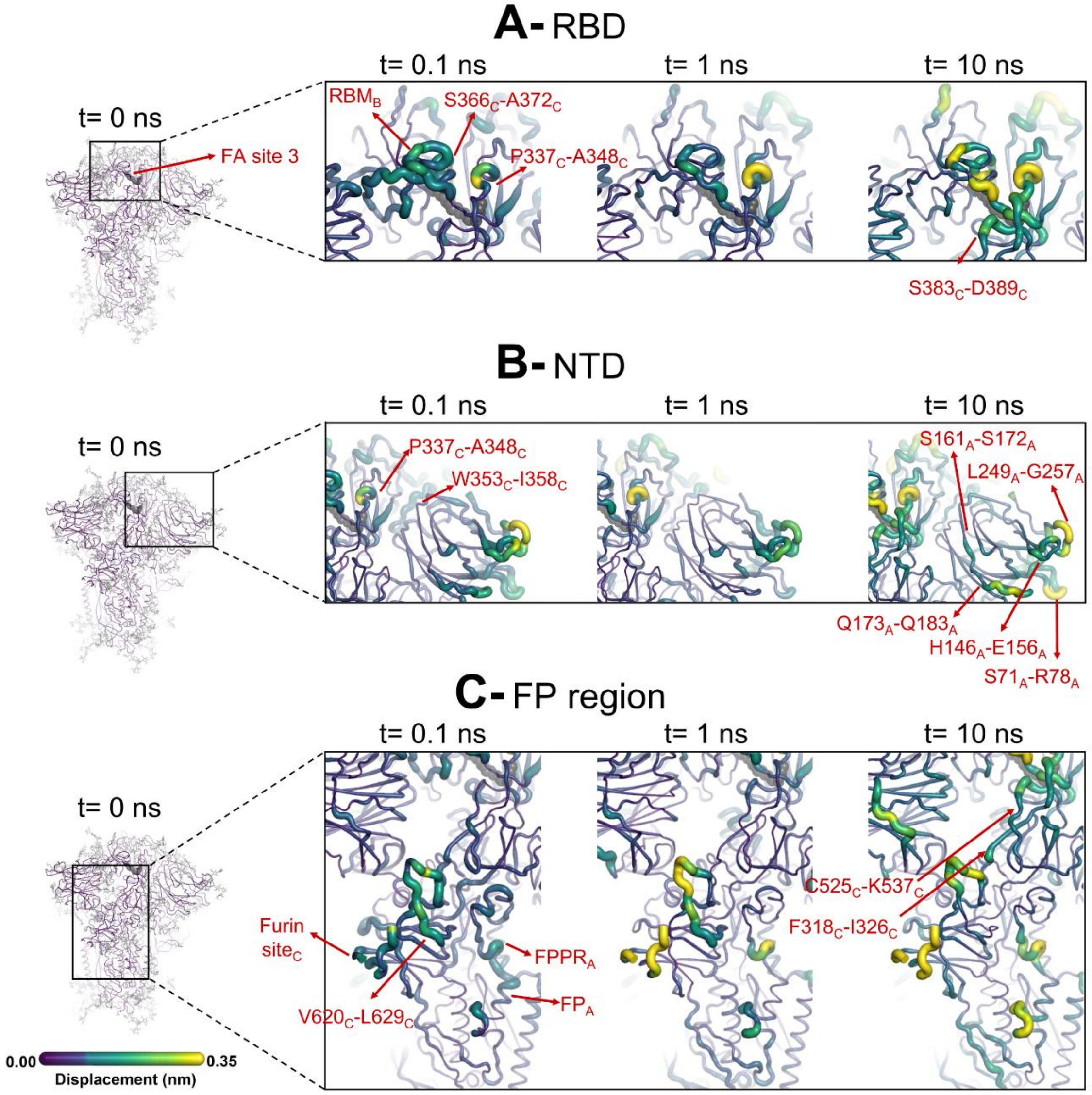


**Figure S15. Evolution of the D-NEMD structural response from the viewpoint of FA site 3** (between chains B and C) to LA removal. Close-up view of the structural response of the $RBD_{BC}$ (**A**), $NTD_A$ (**B**) and $FP_A$ surrounding regions (**C**) to LA removal. Chain ID (A, B or C) is indicated in subscript. The average $C_\alpha$ displacements at *t*=0, 0.1, 1 and 10 ns after LA removal are mapped onto the starting structure for the equilibrium simulations. The structure colours and cartoon thickness indicate the average $C_\alpha$-displacement values. The dark grey spheres highlight the FA binding site. The glycans are omitted from the panels showing the responses at *t*=0.1, 1 and 10 ns.

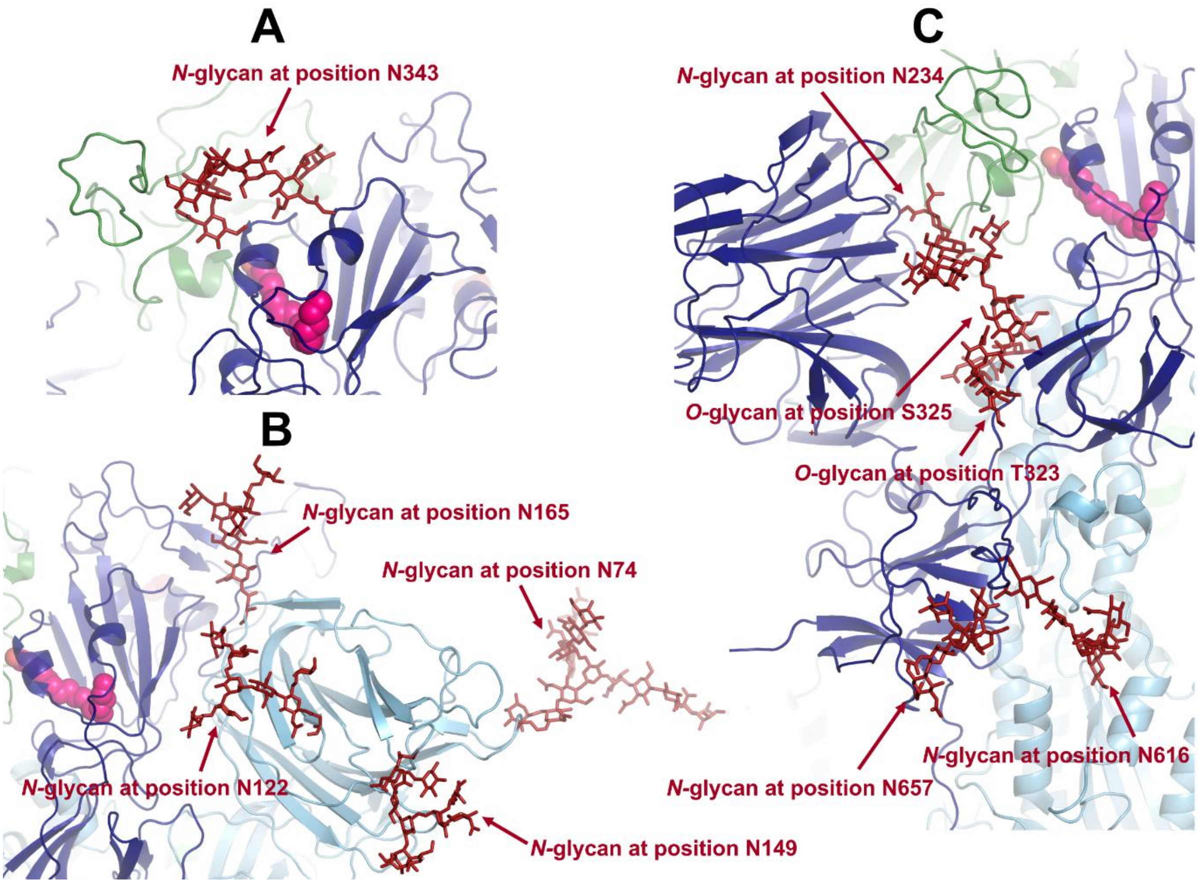


**Figure S16. Example of a conformation showing the position of the glycans located in or close to the allosteric pathways** connecting the FA site to the (**A**) RBM, (**B**) NTD and (**C**) furin cleavage site and FP-surrounding regions (including the FPPR and the S2' cleavage site). Each monomer in the protein is shown in a different colour: dark blue, light blue and green. Glycans are shown with the dark red sticks, and LA molecules are highlighted with magenta spheres.

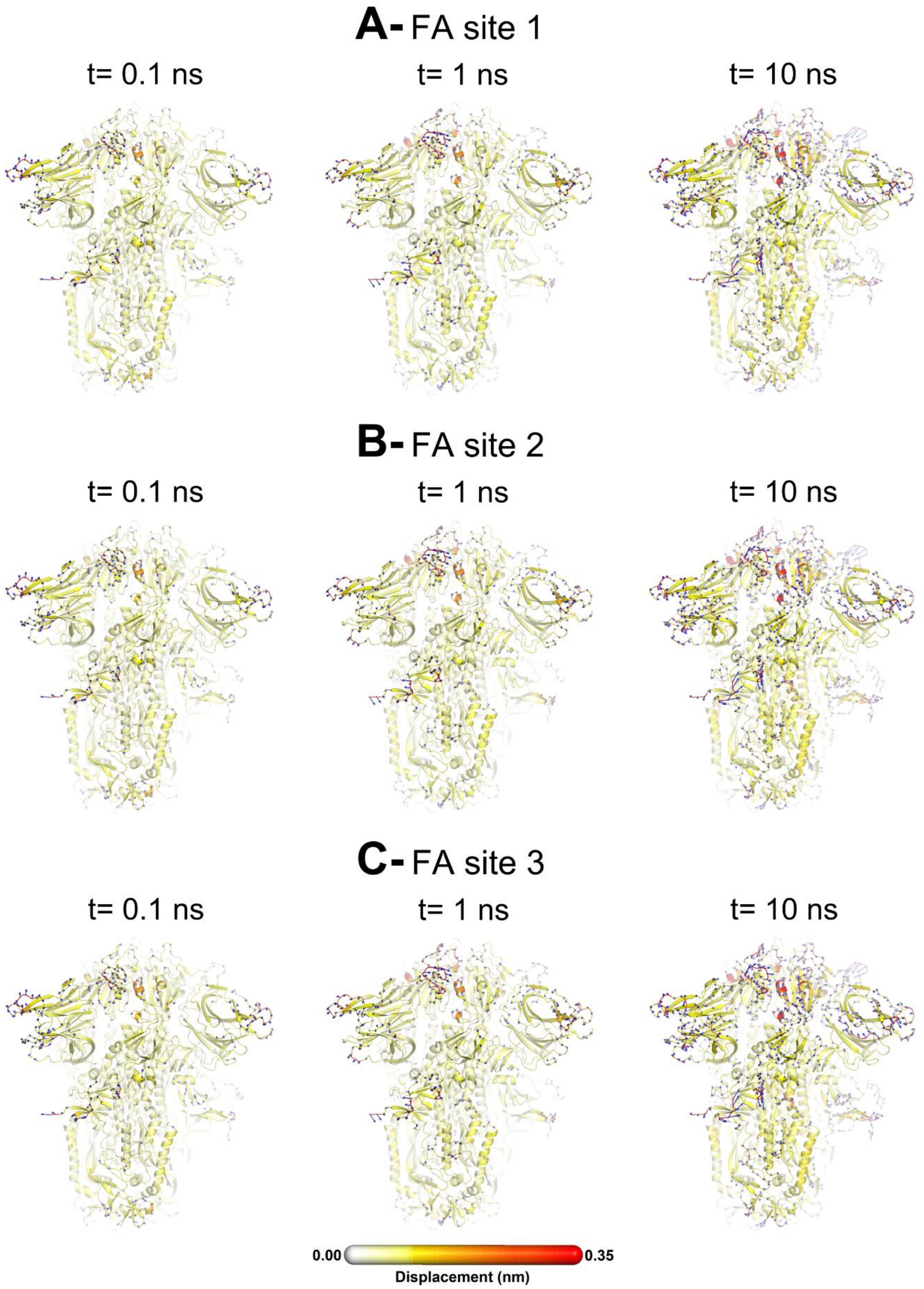


**Figure S17. Average displacement vectors from D-NEMD simulations.** Full view of the protein with the average displacement vectors at time 0.1, 1 and 10 ns. The vectors shown were determined by averaging $C_\alpha$ displacement vectors between the equilibrium and nonequilibrium trajectories over the 210 replicas. Vectors with length ≥0.1 nm are displayed as blue arrows with a scale-up factor of 10. Displacement magnitudes are represented on a white-yellow-orange-red scale. Please zoom in on the image for a detailed visualisation.

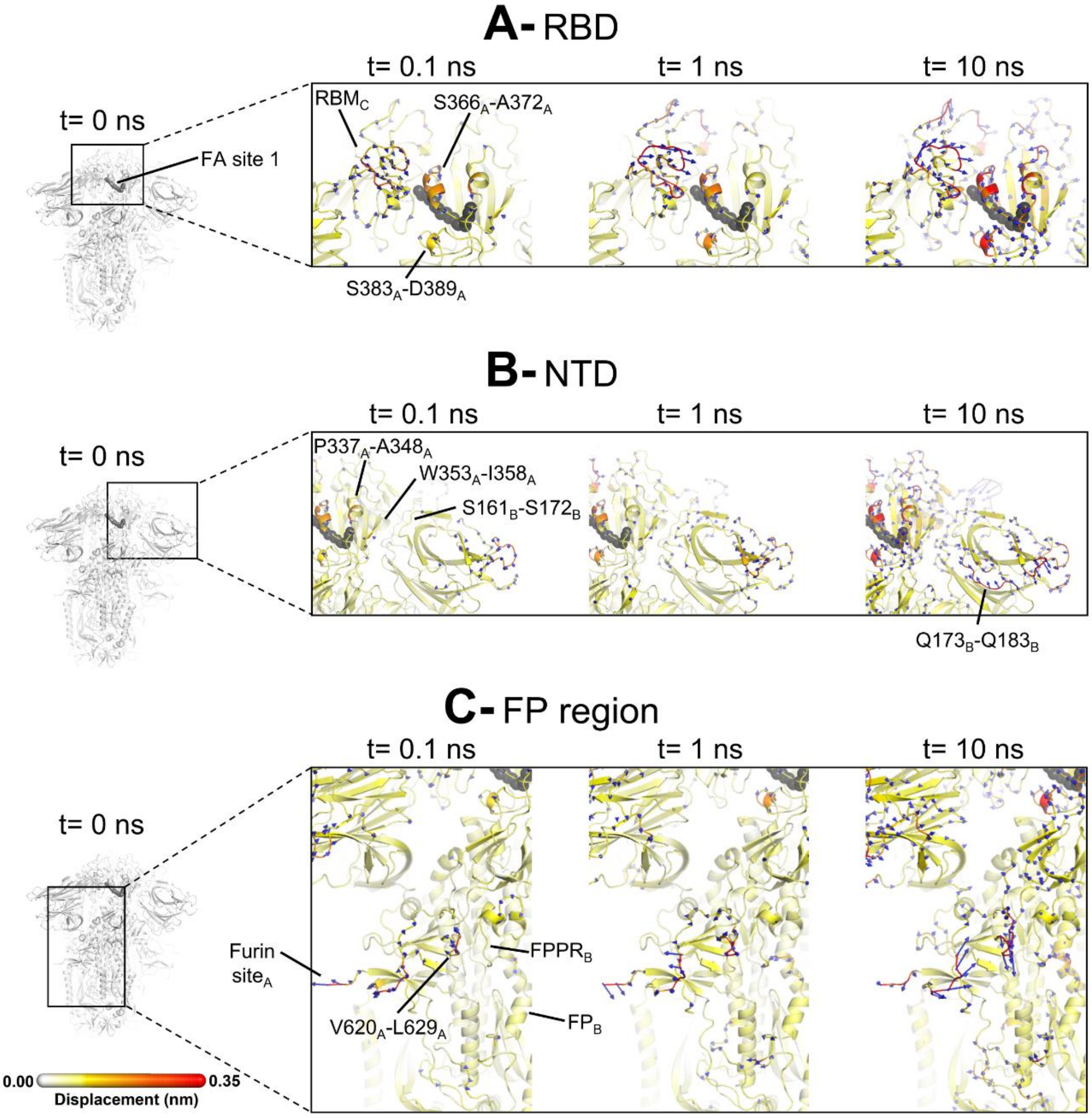


**Figure S18. Direction of the motions from D-NEMD in response to LA removal**. Close-up view of the direction of the responses of the RBD$_{CA}$ (A), NTD$_B$ (B), and FP$_B$ surrounding regions (C). Chain ID (A, B or C) is indicated in subscript. The average $C_\alpha$ displacement vectors at times 0.1, 1 and 10 ns are shown. These vectors were determined by averaging $C_\alpha$ displacement vectors between the equilibrium and nonequilibrium trajectories over the 210 replicas. Vectors with a length ≥0.1 nm are displayed as blue arrows with a scale-up factor of 10. The average displacement magnitudes are represented on a white-yellow-orange-red scale. The dark grey spheres represent the FA site. This figure shows the direction of the responses around FA site 1, which is located at the interface between chains C and A. Please zoom in on the image for a detailed visualisation.

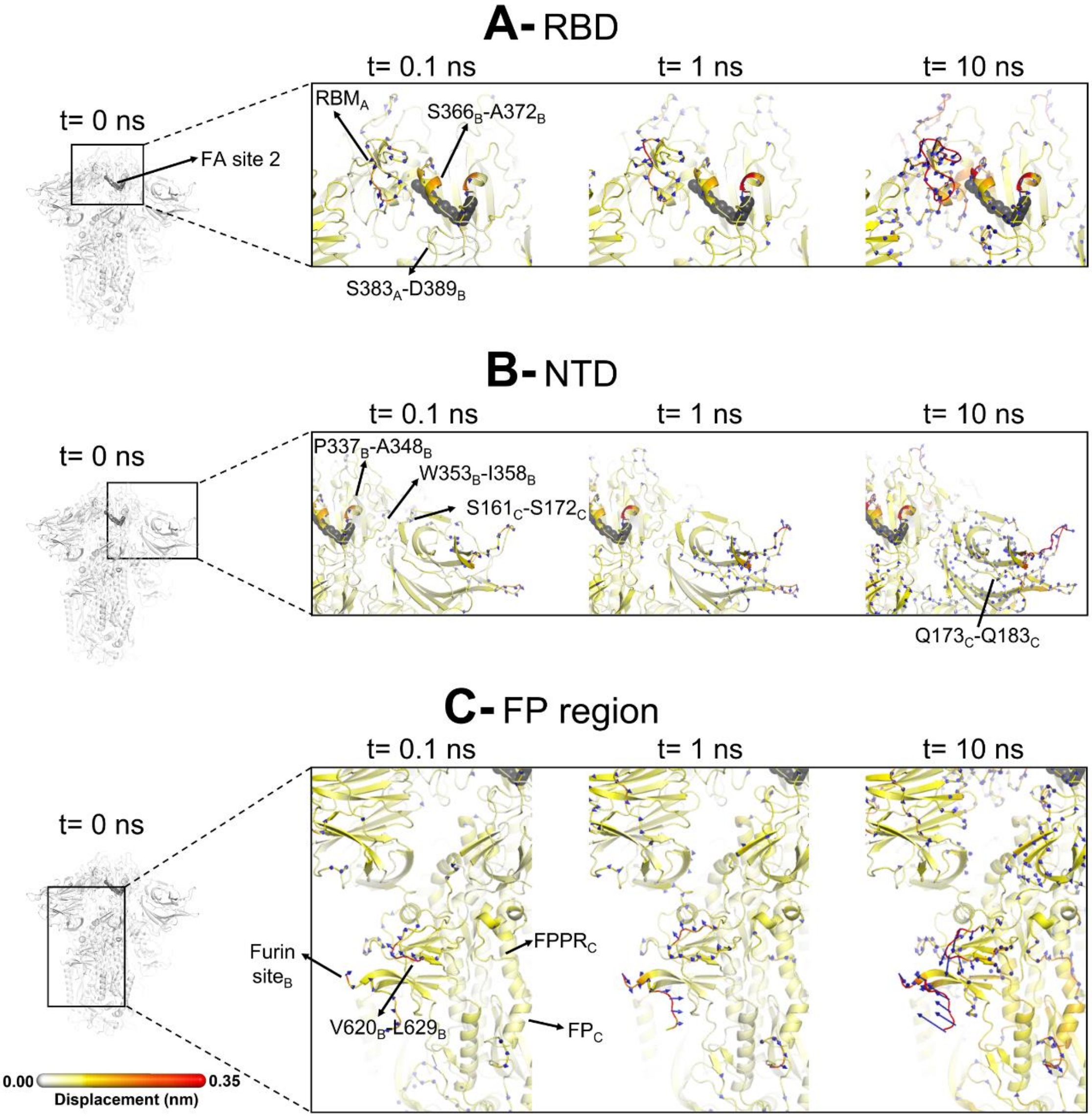


**Figure S19. Direction of the motions from D-NEMD in response to LA removal**. Close-up view of the direction of the responses of the RBD$_{AB}$ (A), NTD$_C$ (B), and FP$_C$ surrounding regions (C). Chain ID (A, B or C) is indicated in subscript. The average $C_\alpha$ displacement vectors at times 0.1, 1 and 10 ns are shown. This figure shows the direction of the responses around FA site 2, which is located at the interface between chains A and B. For more details, see the legend of Figure S17.

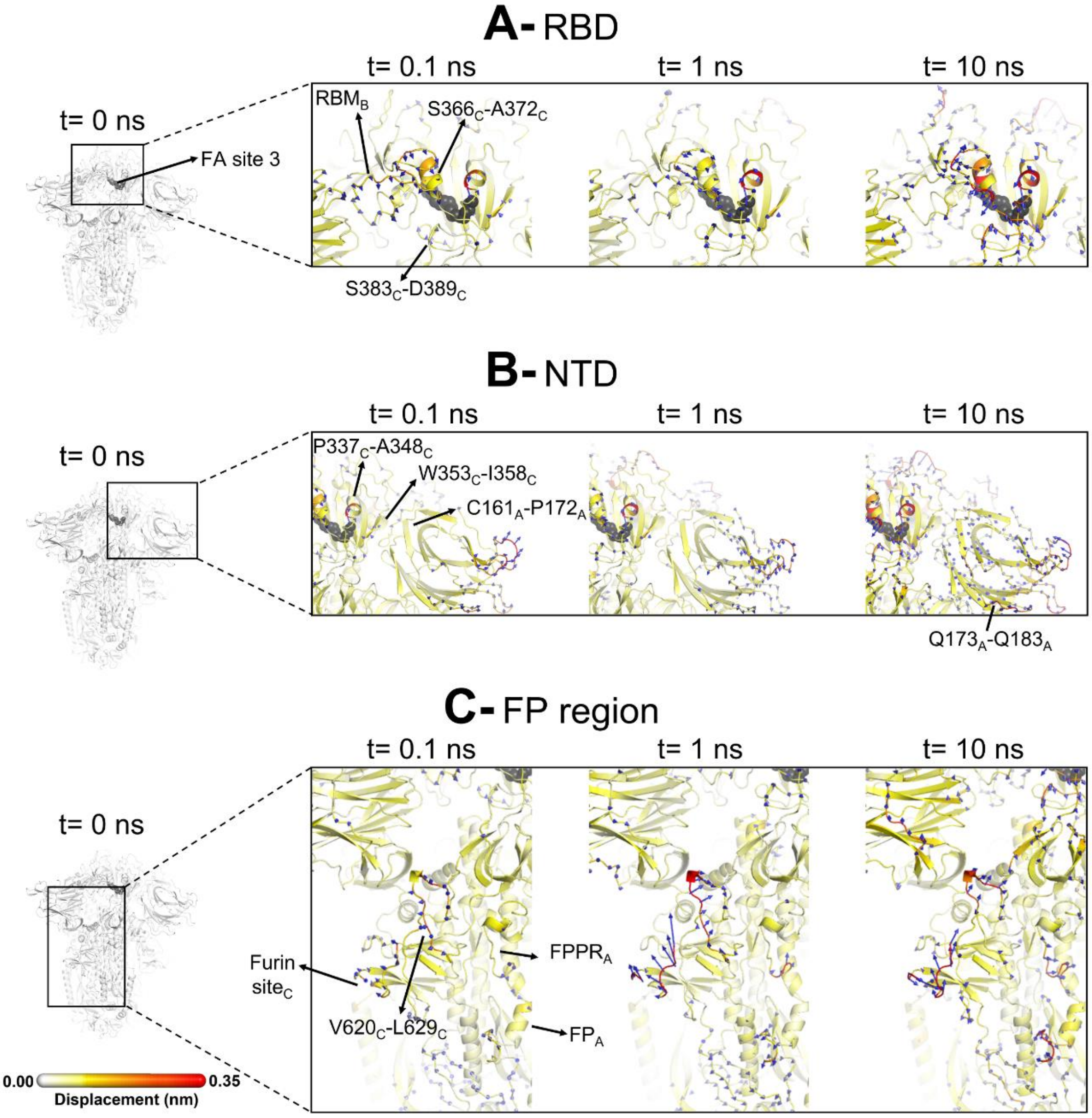


**Figure S20. Direction of the motions from D-NEMD in response to LA removal**. Close-up view of the direction of the responses of the $RBD_{BC}$ (A), $NTD_A$ (B), and $FP_A$ surrounding regions (C). Chain ID (A, B or C) is indicated in subscript. The average $C_\alpha$ displacement vectors at times 0.1, 1 and 10 ns are shown. This figure shows the direction of the responses around FA site 3, which is located at the interface between chains B and C. For more details, see the legend of Figure S17.

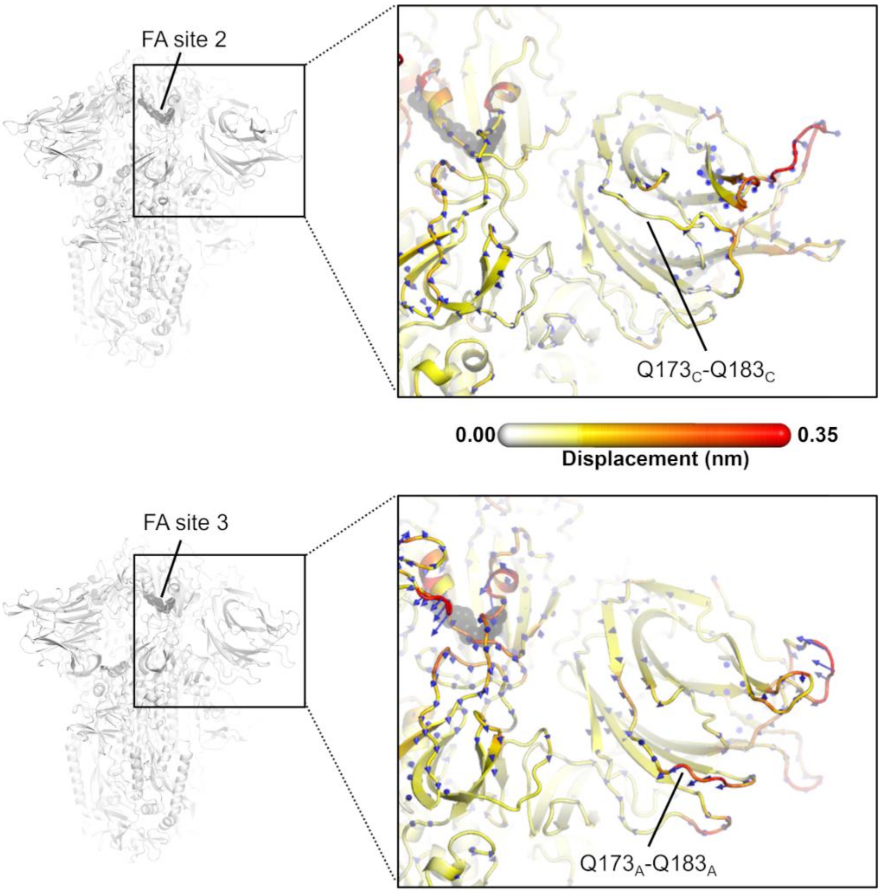


**Figure S21. Direction of the D-NEMD structural responses of $NTD_C$ (top panel) and $NTD_A$ (bottom panel) at *t*=10 ns**. Similarly to the responses of $NTD_B$ (in main text Figure 5A), the Q173-Q183 region in $NTD_A$ (situated closer to FA site 3) shows an outward motion upon LA removal. The magnitudes of the displacements are represented on a white-yellow-orange-red. Vectors with a length $\geq$0.1 nm are displayed as blue arrows with a scale-up factor of 10. The dark grey spheres represent the FA site.

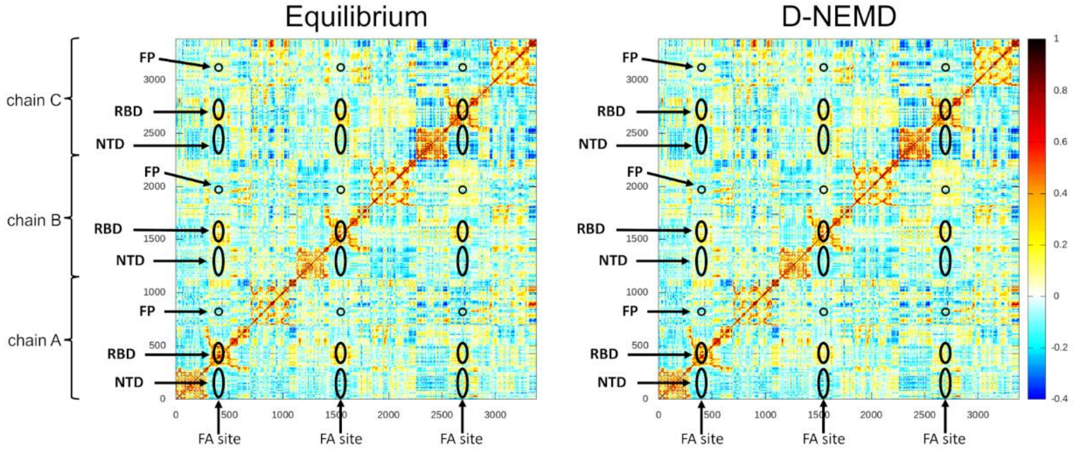


**Figure S22. Cross-correlation maps for the equilibrium (LA-bound) and the D-NEMD simulations.** For the equilibrium map (left panel), the correlations were calculated for all $C_\alpha$ atoms over the three equilibrium trajectories, whereas for the D-NEMD maps (right panel), the correlations were determined using all 210 D-NEMD simulations performed. Atoms that systematically move in the same direction have a correlation value higher than zero, while those moving in opposite directions have a correlation value lower than 0. White regions indicate no correlation. Yellow, orange and red colours indicate low, moderate and significant positive correlations, while cyan and dark blue represent moderate and significant negative correlations. The positions of some key structural features are highlighted in black, namely the N-terminal domain (NTD), receptor-binding domain (RBD), and the fusion peptide (FP). Please zoom in on the image for detailed visualisation.

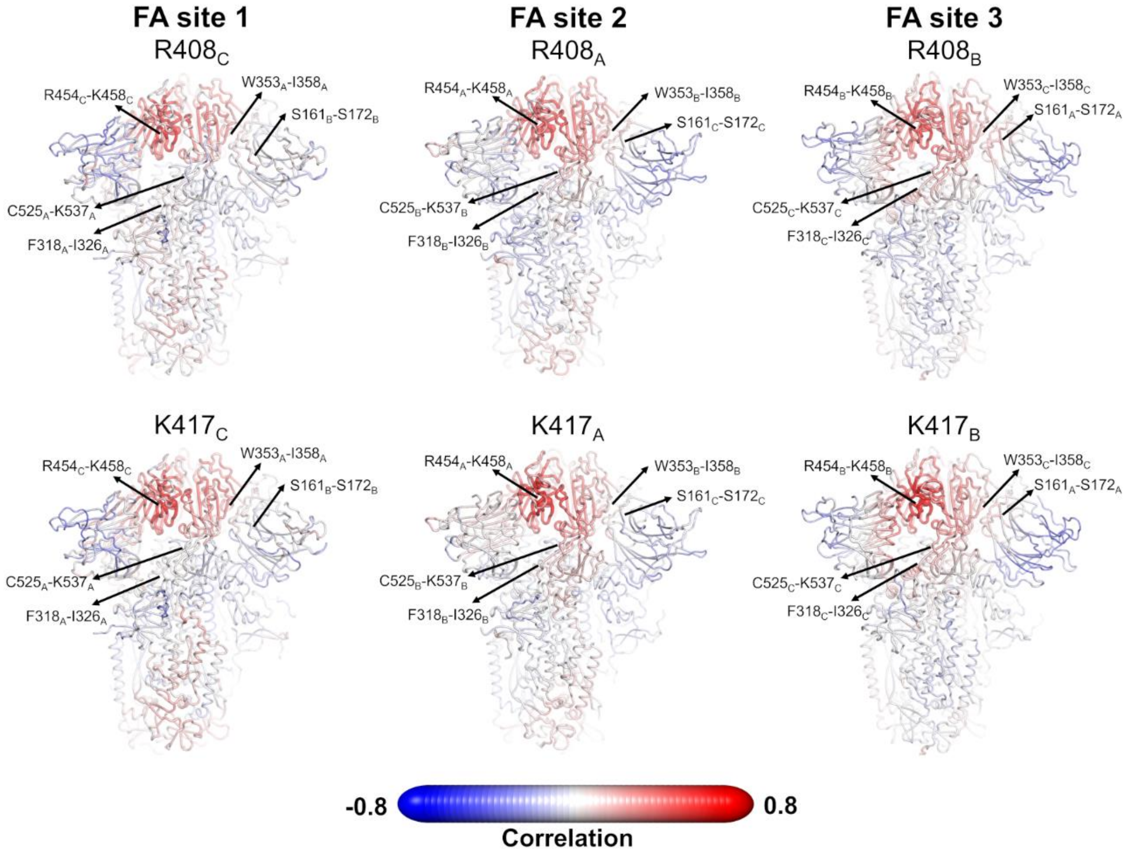


**Figure S23. Pearson correlations for the Cα atom of R408 and K417 from the equilibrium simulations.** R408 and K417 are located in the FA site (see main text Figure 1B) and can directly interact with the carboxylate headgroup of LA. The correlations between the Cα atom of R408 and K417 and all the remaining $C_\alpha$ atoms are shown. Note that similar correlation profiles were observed for R408 and K417 using the D-NEMD trajectories. Atoms that systematically move in the opposite direction to R408 or K417 have strong negative correlation values, whereas those systematically moving in the same direction show strong positive correlations. Atoms whose movements relative to each residue are uncorrelated present a correlation value of 0. The location of R408 (top panel) and K417 (bottom panel) is highlighted with a sphere. Each residue/region is subscripted with its chain ID (A, B or C). Please zoom in on the image for detailed visualisation.

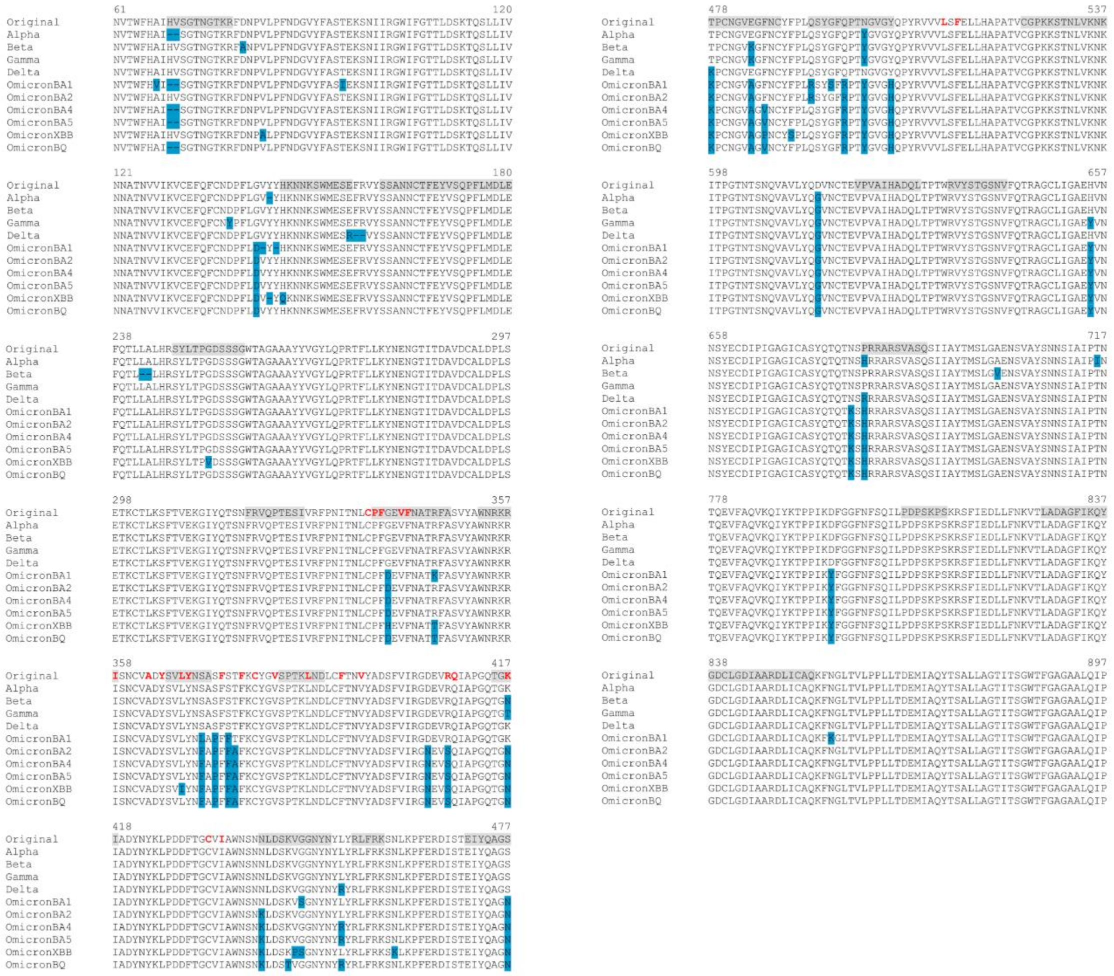

**Figure S24. Sequence alignments for the regions connected to the FA site.** The residues lining the FA site in the original protein are shown in red. The blue boxes represent the location of the substitutions, deletions, and insertions reported for Alpha (SARS-CoV-2 Alpha), Beta (SARS-CoV-2 Beta), Gamma (SARS-CoV-2 Gamma), Delta (SARS-CoV-2 Delta) and several Omicron sub-variants, namely BA.1.1 (SARS-CoV-2 Omicron BA.1.1), BA.2 (SARS-CoV-2 Omicron BA.2), BA.4.1 (SARS-CoV-2 Omicron BA.4.1), BA.5 (SARS-CoV-2 Omicron BA.5), BQ.1.1 (SARS-CoV-2 Omicron BQ.1.1) and XBB.1.5 (SARS-CoV-2 Omicron XBB.1.5) (32). The grey boxes highlight the regions in the original spike shown to respond to the removal of LA from the FA sites. Please zoom in on the image for detailed visualisation.

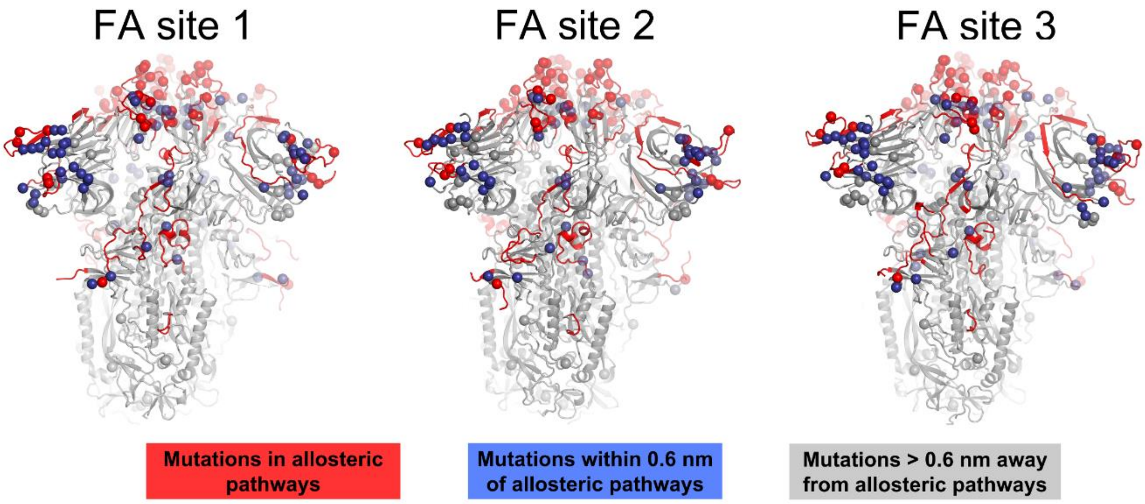


**Figure S25**. **Location of substitutions, deletions and insertions reported for several variants of concern and their relationship to the allosteric pathways identified by D-NEMD here**. The communication pathways captured by D-NEMD (Figure 3 in the main manuscript and Figures S14-S15) are coloured in red, with the mutations lying directly in those pathways shown with red spheres. The mutations closely located to the allosteric pathways (within 0.6 nm of any atom forming the paths) are shown with blue spheres, whereas those further away are shown with grey spheres. The spheres highlight the position of the substitutions, deletions and insertions reported for Alpha (outbreak.info SARS-CoV-2 Alpha), Beta (outbreak.info SARS-CoV-2 Beta), Gamma (outbreak.info SARS-CoV-2 Gamma),Delta (outbreak.info SARS-CoV-2 Delta) and several Omicron sub-variants, namely BA.1.1 (outbreak.info SARS-CoV-2 Omicron BA.1.1), BA.2 (outbreak.info SARS-CoV-2 Omicron BA.2), BA.4.1 (outbreak.info SARS-CoV-2 Omicron BA.4.1), BA.5 (outbreak.info SARS-CoV-2 Omicron BA.5), BQ.1.1 (outbreak.info SARS-CoV-2 Omicron BQ.1.1) and XBB.1.5 (outbreak.info SARS-CoV-2 Omicron XBB.1.5) (32). The red spheres correspond to mutations, deletions and insertions located in the allosteric pathways identified using D-NEMD simulations and the blue spheres correspond to the changes directly contacting these pathways. The grey spheres show changes with no direct contact to the pathways. The protein is shown in light grey, with the allosteric pathways connecting the FA site to important functional regions of the spike coloured in red.

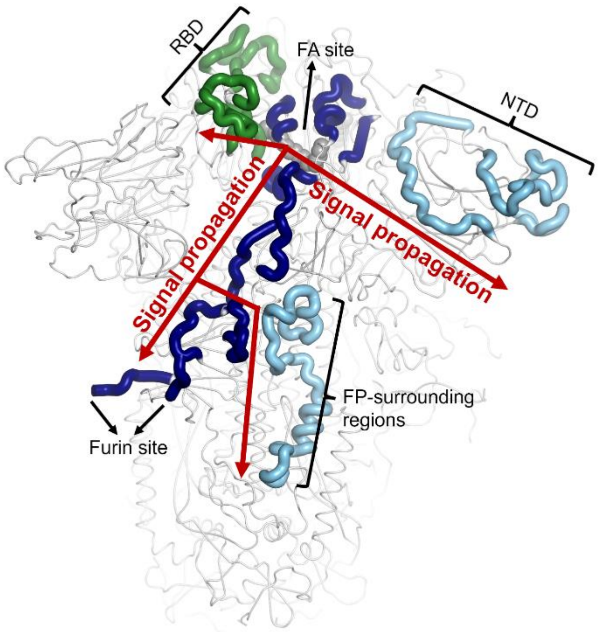

**Figure S26. Scheme representing the pathways identified by D-NEMD connecting the FA site to the RBD, NTD and FP-surrounding regions**. The regions of the pathways belonging to chains A, B and C (which correspond to regions responding to LA removal in Figure 3 in the main manuscript) are coloured in dark blue, light blue and green, respectively. The LA molecule bound to FA site 1 is shown with grey spheres. Note that similar communication pathways are observed from the FA sites 2 and 3.
